# Astrocyte-selective AAV-ADAMTS4 gene therapy combined with hindlimb rehabilitation promotes functional recovery after spinal cord injury

**DOI:** 10.1101/797696

**Authors:** Jarred M. Griffin, Barbara Fackelmeier, Connor A. Clemett, Dahna M. Fong, Alexandre Mouravlev, Deborah Young, Simon J. O’Carroll

**Author notes:** Corresponding author: Dr Jarred Griffin. **Addresses and contact information** Dr Jarred Griffin, Corresponding author, Present address: German Center for Neurodegenerative Diseases (DZNE), Sigmund-Freud-Strasse 27, 53127 Bonn, Germany, Miss Barbara Fackelmeier: The University of Auckland. 85 Park Road, Grafton, New Zealand, Mr Connor Clemett: The University of Auckland. 85 Park Road, Grafton, New Zealand, Dr Dahna Fong: The University of Auckland. 85 Park Road, Grafton, New Zealand, Dr Alexandre Mouravlev: The University of Auckland. 85 Park Road, Grafton, New Zealand, Associate Professor Deborah Young: The University of Auckland. 85 Park Road, Grafton, New Zealand, Dr Simon O’Carroll: The University of Auckland. 85 Park Road, Grafton, New Zealand.

## Abstract

Chondroitin sulphate proteoglycans (CSPGs) are inhibitors to axon regeneration and plasticity. Bacterial chondroitinase ABC degrades CSPGs and has been extensively reported to be therapeutic after SCI but there remain concerns for its clinical translation. A disintegrin and metalloproteinase with thrombospondin motifs-4 (ADAMTS4) is a human enzyme that catalyses the proteolysis of CSPG protein cores. Infusion of ADAMTS4 into the damaged spinal cord was previously shown to improve functional recovery after SCI, however, this therapy is limited in its enzyme form. Adeno-associated viral (AAV) vector gene therapy has emerged as the vector of choice for safe, robust and long-term transgene expression in the central nervous system. Here, an AAV expression cassette containing ADAMTS4 under the control of the astrocytic GfaABC_1_D promoter was packaged into an AAV5 vector. Sustained expression of ADAMTS4 was achieved *in vitro* and *in vivo*, leading to widespread degradation of CSPGs. AAV-ADAMTS4 resulted in significantly decreased lesion size, increased sprouting of hindlimb corticospinal tract axons, increased serotonergic fiber density caudal to the injury, and improved functional recovery after moderate contusive SCI. Hindlimb-specific exercise rehabilitation was used to drive neuroplasticity towards improving functional connections. The combination of hindlimb rehabilitation with AAV-ADAMTS4 led to enhanced functional recovery after SCI. Thus, widespread and long-term degradation of CSPGs through AAV-ADAMTS4 gene therapy in a combinational approach with rehabilitation represents a promising candidate for further preclinical development.

## Introduction

Axons In the adult mammalian central nervous system (CNS) have limited regenerative capacity following spinal cord injury (SCI), leading to permanent loss of sensory and motor functions in affected individuals. This is due to both poor intrinsic regenerative capacity of adult neurons as well as extrinsic inhibitory factors present in the extracellular matrix such as chondroitin sulphate proteoglycans (CSPGs) and myelin-associated molecules (Curcio and Bradke, 2018; Vogelaar, 2016). CSPGs are well established to be potent inhibitors of axonal growth and plasticity *in vitro* and *in vivo,* which is believed to be exerted through their negatively charged, covalently attached, chondroitin-sulphate glycosaminoglycan side chains (CS-GAGs; (Rudge and Silver, 1990; Snow et al., 1996; Snow et al., 1990; Snow and Letourneau, 1992). Degradation of the CS-GAGs by the bacterial enzyme chondroitinase ABC (ChABC) has been shown in a variety of models to remove the inhibitory influence exerted by these molecules (Bradbury and Carter, 2011). When infused into the contused rat spinal cord, ChABC increased sprouting of corticospinal tract (CST) fibres, restored functional connections to produce a modest improvement in motor function as assessed by performance in beam walking and grid walking tasks (Bradbury et al., 2002; Garcia-Alias et al., 2009). More recently, a mammalian-compatible ChABC (mChABC) gene was engineered to address the possible risk for clinical use of the bacterial enzyme and to allow for expression from mammalian cells (Muir et al., 2010). Lentiviral vector (LV) gene delivery of mChABC achieved long-term expression in the contused rat spinal cord which resulted in large-scale CSPG degradation and improved behavioural scores (Bartus et al., 2014). This team further developed an immune-evasive Tet-on inducible LV-mChABC vector that allows for regulated gene therapy in efforts to move towards clinical translation (Burnside et al., 2018). While LV-mChABC gene therapy has undeniable benefit, there remains concern about the safety of the use of the therapy in humans. Lentiviral vectors risk insertional mutagenesis, transgene oncogenesis, generation of replication-competent retrovirus’s (Schlimgen et al., 2016), as well as the possibility of the modified prokaryote ChABC enzyme still being capable of eliciting an immune response. Is there a mammalian endogenous alternative? ADAMTSs (a disintegrin and metalloproteinase with thrombospondin motifs) may represent such an alternative. ADAMTSs are mammalian endogenous enzymes that catalyse the proteolysis of CSPG core proteins (Gottschall and Howell, 2015). The role of ADAMTSs in the injured spinal cord is relatively unexplored. They are expressed through the CNS and ADAMTS4 appears to be the most highly expressed in the brain (Yuan et al., 2002). When infused into the severely contused rat spinal cord for 14 days, ADAMTS4 treated rats had a significant improvement in BBB (Basso, Beattie and Bresnahan) locomotor scores, equalling the benefit of ChABC infusion (Tauchi et al., 2012). A gene therapy approach utilising an adeno-associated viral vector (AAV) and expressing human ADAMTS4 may be an advantageous alternative to LV-mChABC. AAV vectors have emerged as the vector of choice for safe, robust and long-term transgene expression in the CNS with their well-established clinical safety profile (Naso et al., 2017). Our lab previously developed an astrocyte-targeting AAV vector for spinal cord gene therapy applications (Griffin et al., 2019). This vector contains a truncated GFAP promoter (GfaABC_1_D) packaged into an AAV5 serotype and is capable of transducing a large volume of spinal cord tissue with selective transgene expression in astrocytes without affecting behavioural outcomes. It is becoming increasingly clear that astrocytes play a critical role in SCI pathogenesis as well as other neurotrauma or neurodegenerative conditions. Therefore, AAV vectors that exhibit a propensity toward transducing astroglial cell populations are attractive candidates and could potentially be more beneficial than gene therapies that typically display neuron-centric transduction profiles.

It is also becoming apparent that using appropriate rehabilitative strategies can help direct the strengthening of correct connections following regenerative therapies (Fawcett and Curt, 2009; Loy and Bareyre, 2019). How exactly rehabilitation influences plasticity is not well understood, however, appropriate rehabilitation regimes can prove to be astonishingly effective. For example, task-specific rehabilitation combined with ChABC almost completely restored some motor tasks (Chen et al., 2017; Garcia-Alias et al., 2009). Furthermore, specifically training the hindlimbs using bipedal treadmill exercise after thoracic spinal cord injury significantly improved several parameters of motor functions when combined with the plasticity-promoting treatment, anti-Nogo-A antibody (Chen et al., 2017). Rehabilitation may enhance the ADAMTS4-induced improvements to behavioural scores that were shown by Tauchi and colleagues.

Here we investigated the therapeutic potential of an astrocyte-targeting AAV vector expressing human ADAMTS4 in a rodent spinal cord contusion model. We observed that this treatment significantly decreased lesion size, increased sprouting of hindlimb corticospinal tract axons, increased serotonergic fiber density caudal to the injury, and improved functional recovery after moderate contusive SCI. We further show that the combination of hindlimb rehabilitation further enhanced functional recovery.

## Materials and methods

### Vector construction and packaging

Polymerase chain reaction was used to expand ADAMTS4 cDNA to be cloned into a suitable plasmid cassette. A plasmid containing human ADAMTS4 cDNA (Origene: RC09226; NM_005099) was used as a template and amplified using a ThermoFisher Accuprime PCR kit as per the manufacturer’s instructions and the forward primer *5’ TAT TAT ACT AGT GCT GCA GTA CCA GTG CCA TG 3’* and the reverse primer *5’ CTA GTA AAG CTT CGG GAT AGT GAG GTT ATT TC 3’* (Fig. S1 a) The ADAMTS4 PCR product was then purified and cloned into an AAV expression plasmid; the ADAMTS4 transgene was followed by a woodchuck hepatitis virus post-transcriptional regulatory element (WPRE) and bovine growth hormone polyadenylation signal (BGHpA), flanked by AAV2 inverted terminal repeats. DNA sequences were confirmed by restriction digest analysis (Fig. S1 a, b) and DNA sequencing (Massey Genome Service; Massey University, New Zealand) and the following expression cassette was generated in this study: pAM/ITR-GfaABC_1_D-ADAMTS4-WPRE-BGHpA-ITR (termed AAVA-ADAMTS4; Fig. S2). Expression cassettes were packaged into the AAV5 serotype and purified as previously described (Mudannayake et al., 2016). The genomic titre of the vector was determined as previously described with primers designed to WPRE and titre-matched to our control vector: pAM/ITR-GfaABC_1_D-dYFP-WPRE-BGHpA-ITR (termed AAV-dYFP) (During et al., 2003). The purity of the vector was determined using SDS-PAGE for viral capsid proteins the and the absence of non-specific bands across all vectors indicates the vector stocks are of high purity (supplementary information; Fig. S1 c).

### Primary astrocyte cell culture

Animal procedures were approved by the University of Auckland Animal Ethics Committee and performed in accordance with the New Zealand Animal Welfare Act 1999. Primary spinal cord astrocytes were isolated from Sprague-Dawley rat pup postnatal days 0-2. The spinal cords of 2 pups (per culture) were excised and astrocytes were harvested as previously described (Kerstetter and Miller, 2012). Cultures were maintained for at least 4 weeks in growth media (DMEM supplemented with 10% fetal bovine serum (FBS), penicillin-streptomycin-neomycin (PSN), 1x Fungizone, and 1x Glutamax) with media changes every 3-4 days before plating for experimentation. Cells were plated into 96 well plates at a density of 4.0×10^5^ cells/well. Two days later, cultures were transduced with AAV-ADAMTS4 or AAV-dYFP (4×10^9^ vg or 8×10^9^ vg), and stimulated with 10 ng/mL TGFβ1 48 hours later. After three days, the cultures were fixed with 4% paraformaldehyde in phosphate buffered saline (PFA-PBS). Cultures were repeated a further four times (n = 4 biological repeats; 8 pups used; 3 technical repeats within each experiment and three images for each technical repeat).

### Animals, surgical procedure and AAV injections

Adult female Sprague Dawley rats weighing 200-300g were used in this study (obtained from the Vernon Jansen Unit, University of Auckland, New Zealand). An acute study ending 4 weeks after SCI and another ending 13 weeks after SCI were conducted (Fig. 1). The acute study was conducted to assess whether a control reporter AAV vector (contusion + AAV-dYFP; *n* = 9), or the therapeutic AAV-ADAMTS4 vector (contusion + AAV-ADAMTS4; *n* = 9) affected histological and behavioural results compared to no treatment (contusion only; *n* = 9). Following this, a long term study was carried out to investigate the therapeutic potential of AAV-ADAMTS4 and the influence of hindlimb rehabilitation (Fig. 1). There were five groups in this study: contusion only (*n* = 14), contusion + AAV-ADAMTS4 (*n* = 13), contusion + rehabilitation (*n* = 12), contusion + AAV -ADAMTS4 + rehabilitation (*n* = 9), and sham surgery + AAV-ADAMTS4 (*n* = 4). This sample size is consistent with our previous study (Griffin et al., 2019) and generated using a type 1 error threshold (α) ≤ 0.05 and power (1 β) ≥ 0.80.

**Figure 1:**
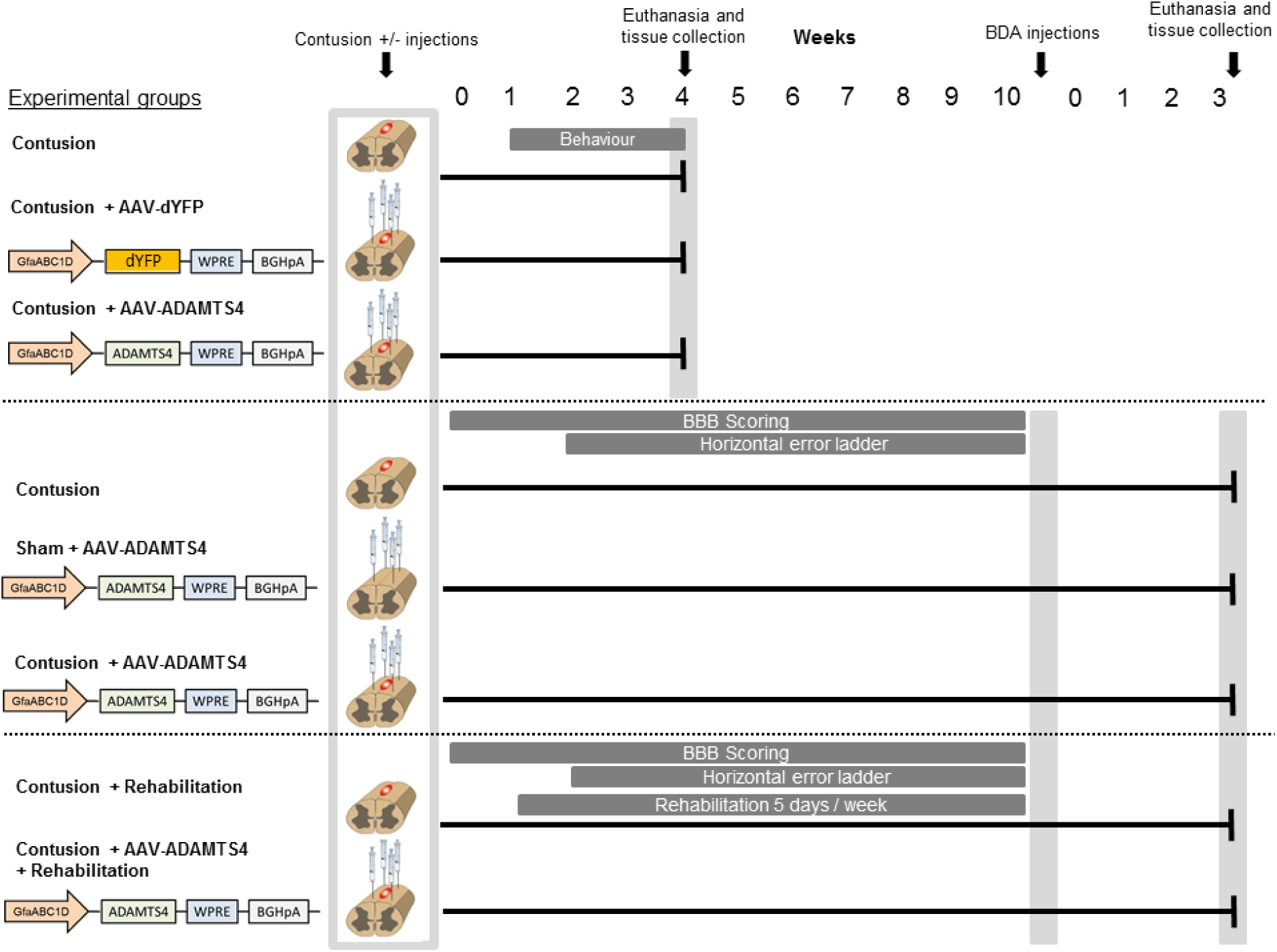
A schematic diagram of the study design. An acute study ending 4 weeks after SCI and another ending 13 weeks after SCI were conducted. To investigate transgene expression and toxicity resulting from the vector, animals were randomly assigned to three treatment groups and tissue was collected 4 weeks following 175 KDyne contusion injuries. To investigate the therapeutic benefit of AAV-ADAMTS4 alone or in combination with hindlimb rehabilitation animals were randomly assigned to experimental groups and on day 0 received a 175 KDyne contusion injury at T10. Those in treatment groups received four injections of AAV-ADAMST4 at each corner of the injury site. Those assigned to the rehabilitation groups were subjected to hindlimb rehabilitation one week after the injury. At week 10, animals received injections of BDA into the hindlimb motor cortex for tracing of hindlimb CST axons. Three weeks later, the animals were euthanised and their spinal cords and brains were collected for histological analyses. This figure was produced using Servier Medical Art, available from www.servier.com/Powerpoint-image-bank. Servier Medical Art by Servier is licensed under a Creative Commons Attribution 3.

The animals were randomly assigned to experimental groups prior to surgery. For surgery, animals were anaesthetised with isoflurane and given subcutaneous injections of local anaesthetic at the surgery site (Bupivicane; 1 mg/kg), analgesic (Carprofen; 5 mg/kg) and antibiotic (Enrofloxacin; 5 mg/kg). A laminectomy was performed to expose the spinal cord at spinal level T10 and impacted in the midline with a force of 175 KDyne using an Infinite Horizon Impactor (Precision Systems Instrumentation). Data recorded from the IH impactor revealed that the actual forces delivered to the cords were within 10% of the intended 175 KDyne. Immediately after contusion injury, the rats were transferred to a stereotaxic frame (David Kopf Instruments). A 10 µL Hamilton Neuros syringe coupled to a Micro Syringe Pump Controller (World Precision Instruments) was used to inject viral vectors into the spinal cord (4×10^9^ vg; 1 µL) at a depth of 1 mm was made at each corner of the lesion site (four in total) at a rate of 200 nL/min. Rats were housed one per cage after surgery for two weeks and then housed four per cage in a temperature and humidity controlled atmosphere with a 12 hour light/dark cycle and *ad libitum* access to food and water. Animals were administered subcutaneously antibiotics (Enrofloxacin; 5 mg/kg), analgesic (Carprofen; 5 mg/kg, Buprenorphine; 0.05mg/kg), and supplementary fluid (0.9% saline, 3 mL) for three days. Bladders will be expressed manually three times daily until the normal voiding response returned.

### Hindlimb rehabilitation

For animals assigned to rehabilitation groups, training began one week after surgery using a five-lane rodent treadmill (Pan Lab/Harvard Apparatus, #LE8710RTS). A custom built frame was constructed and rats were suspended in handmade, custom fitted jackets so that only their hindlimbs were in contact with the treadmill belt. This apparatus allowed for animals to be kept upright whilst having weight bearing movements on their hindlimbs. For the first week of the rehabilitation program, rats were trained for 10 minutes a day at a speed of 5 cm/sec. From weeks 2-10, the rats were then trained for 30 minutes a day, five days a week, at a speed of 25-35 cm/sec.

### BDA anterograde tracing of hindlimb corticospinal tract axons

Ten weeks after spinal cord injury surgery, biotinylated dextran amine (BDA) was injected into the hind-limb motor cortex of animals. Rats were anesthetised by isoflurane and their scalp was shaved and local analgesia (Bupivicane 1mg/kg) was injected subcutaneously. The animal was placed in a stereotaxic frame (David Kopf Instruments) and the skull exposed. Four bilateral burr holes were made at following coordinates in reference to Bregma: 1) 0.5 mm posterior, 1.5 mm lateral; 2) 2.0 mm posterior, 1.0 mm lateral; 3) 0.5 mm posterior, 3.0 mm lateral; 4) 2.0 mm posterior, 3.0 mm lateral. These coordinates are known to be the location of the hindlimb motor cortex in rats (Bareyre et al., 2004; Neafsey et al., 1986). All injections were made 3 mm from the surface of the skull, and 1 µL of BDA (10% w/v in sterile PBS) injected into each site via a 26 gauge Hamilton syringe needle attached to a micro-infusion pump (World Precision Instruments) at a rate of 200 nL/min.

### Behavioural testing

All behavioural testing was completed by blinded observers. The Basso, Beattie, and Bresnahan (BBB), horizontal error ladder and CatWalk Gait XT were used to test hind limb motor functions. BBB testing was carried out by two blinded observers in a circular open field for 5 minutes each as previously described (Basso et al., 1995). Animals were acclimatised to the open field for three days before testing and testing began the day after surgery (Metz and Whishaw, 2009). For the error ladder test, a 1-meter long ladder with irregularly spaced rungs was used. Animals were acclimatised to the ladder for three days before testing and testing began the day after surgery. Each animal completed three runs of the ladder which was recorded using a GoPro Hero4 camera. The footage was analysed frame-by-frame by a blinded observer to observe footfalls and the total number of errors was recorded over the three runs. For the CatWalk Gait analysis, the parameters of body speed, print width, print area, and stride length were measured and the average of 5 runs is presented.

### Tissue processing

Three weeks following BDA injections the animals were euthanized and their brain and spinal cord were collected for histological analysis. Animals were overdosed with sodium pentobarbitone (Pentobarb; Euthatal 300; 100 mg/kg, i.p) and transcardially perfused with 0.9% saline followed by 4% paraformaldehyde (PFA) in 0.1 M phosphate buffer (PB). Immediately after perfusion, approximately 5 centimetres of the spinal cord with the lesion in the centre and the brain of the animal were removed. The cord and brain were post-fixed in 4% PFA in 0.1 M PB for 4 hours at 4°C before being cryoprotected in 20% and then 30% sucrose in 0.1 M PB. Cryosectioning was performed using a Leica CM3050 S Cryostat. The tissue was frozen inside the cryostat and 1.2 centimetres of the cord with the lesion in the centre was cut. The cord was then embedded in OCT cut onto positively charged microscope slides so that each adjacent section on a slide represented 600 µm of distance. All histological staining, image acquisition and analysis was performed by blinded researchers.

### Luxol fast blue and eosin staining

Luxol fast blue and eosin staining was used to visualise grey and white matter and lesion size in the spinal cord tissue sections. The sections were incubated at in 0.1% Luxol fast blue at 60°C for six hours then differentiated in 0.1% lithium carbonate for 30 seconds before changing to 70% ethanol for 30 seconds and then rinsed off in water. The sections were counterstained with eosin for 30 seconds before being cleared with xylene and mounted. A Leica DMR upright microscope was used to capture 2.5x magnification images of the stained sections. ImageJ software was used to trace the regions of white matter, grey matter and lesion size for every section. This staining was performed twice and the values were averaged to minimise variation in the data.

### Visualisation of BDA-labelled collaterals

Diaminobenzidine (DAB) immunohistochemistry was used to visualise BDA within traced neurons. For BDA detection: sections were first washed with PBST for 20 minutes before being incubated overnight at 4°C with Avidin-Biotin-Peroxidase-Complex reagent (ABC) diluted in PBS + 1% bovine serum albumin (BSA). The sections were then washed in PBS and incubated with biotinyl-tyramide-xx signal amplification reagent (ThermoFisher), as per the manufacturer’s instructions, for two hours at room temperature followed by incubation with ExtrAvidin Peroxidase (Sigma) diluted 1:250 in PBS for two hours at room temperature. Sections were stained with DAB; 0.2 mg/mL DAB, 0.01% hydrogen peroxide, 0.1 M PB pH 7.2. Slides were mounted and imaged using a Leica DMR microscope at 2.5x magnification to image whole sections or at 10x or 40x magnification to obtain higher magnification images. For analysis, in sections 600-1200 µm rostral to the lesion edge, all BDA-labelled axons present outside of the CST bundle were traced in ImageJ and represented as the sum of axons traced (µm).

### Visualisation of ADAMTS4-proteolysis

For ARGxx detection, the same protocol as above was used with the following changes: The primary antibody was mouse anti-ARGxx (Abcam; ab3773) diluted at 1:200 in PBS + 0.01% Triton X-100 (PBST). A biotinylated goat-anti-mouse secondary antibody (Sigma; SAB46000004) was diluted at 1:250 in PBST and then incubated at room temperature for two hours with ABC reagent diluted in PBS + 1% BSA. DAB colour development was performed as above. Stained sections were then imaged using a Leica DMR microscope at 2.5x, 10x and 40x magnification.

### Fluorescent immunohistochemistry and intensity analyses

For fluorescent immunohistochemistry, spinal cord sections or cells were permeabilised using PBST and then incubated overnight at room temperature in primary antibody diluted in PBS-T + 4% goat or donkey serum (rabbit anti-ADAMTS4 Abcam 1:1000; mouse anit-GFAP-Cy3 Sigma 1:500; biotinylated wisteria lectin floribunda WFA, Sigma 1:200; mouse anti-chondroitin sulphate CS56, Abcam 1:200; goat anti-5HT, Abcam 1:500). After two washes, the cells or sections were then incubated with fluorescent-secondary antibodies (Donkey-anti-rabbit/goat/mouse Alexa Fluor 488/594, ThermoFisher 1:500; Streptavidin Alexa Fluor488, Sigma 1:200) at room temperature for two hours. Images were captured using an Olympus FV1000 confocal microscope or with a Nikon Eclipse TE2000-U. For fluorescent intensity analyses, images were thresholded to correct for background and the mean pixel intensities were measured using ImageJ. The same threshold was maintained for all immunostaining groups. For 5HT analysis specifically, the fluorescent intensity from the ventral horns of three sections: 600, 1200 and 1800 µm caudal to the injury measured using ImageJ. The pixel intensity measured at the ventral horns were subtracted from the pixel measured at the dorsal horn within each image.

### Data exclusions and statistical analyses

All behavioural testing, histological staining, image acquisition and analysis were performed by blinded researchers. For the 13 week study, a number of animals were excluded from the study. The criteria for this was if their combined hindlimb BBB score was greater than five at day one after spinal cord surgery, if their BBB scores did not improve past a combined score of eight by the end of the four weeks, and if the animal was euthanized because of complications of the surgery. Furthermore, animals assigned to AAV-ADAMST4 group were excluded if ADAMTS4 expression was not observed within the tissue, indicating a failed injection. This resulted in n = 14 for contusion, n = 13 for contusion + AAV-ADAMTS4, n = 12 for contusion + rehabilitation, n = 9 for contusion + AAV-ADAMTS4 + rehabilitation. Values are expressed as the mean ± standard error of the mean (SEM). Statistical analyses were conducted using Prism version 7.0 (GraphPad, La Jolla, USA). For cell culture experiments and 5HT analysis, one-way ANOVA’s followed by a Tukey’s multiple comparisons was used to determine statistically significant differences between the means of groups split by one independent variable where \**P* < 0.05, \*\**P* < 0.001, \*\*\**P* < 0.0001. For LFB histology and behavioural data, Two-way ANOVA’s followed by a Bonferroni multiple comparisons test were used to compare the contribution to the variance of the variables, and the mean differences in data between groups that have been split on two independent variables where \**P* < 0.05, \*\**P* < 0.001, \*\*\**P* < 0.0001.

## Results

### AAV-dYFP results in robust expression and does not affect astrogliosis or behavioural parameters

We have previously created an astrocyte-targeting AAV5 vector with a truncated GFAP promotor (GfaABC_1_D) to selectively mediate transgene expression within spinal cord for applications in SCI gene therapy (Griffin et al., 2019). This vector robustly transduces primary spinal astrocyte cultures, saturating from 2×10^9^ viral genomes per well (Fig. 2 a, b). Transduction did not affect astrogliosis, as assessed by GFAP levels, nor did transduction alter cell number compared to non-transduced cells (Fig. 2 c, d).

**Figure 2:**
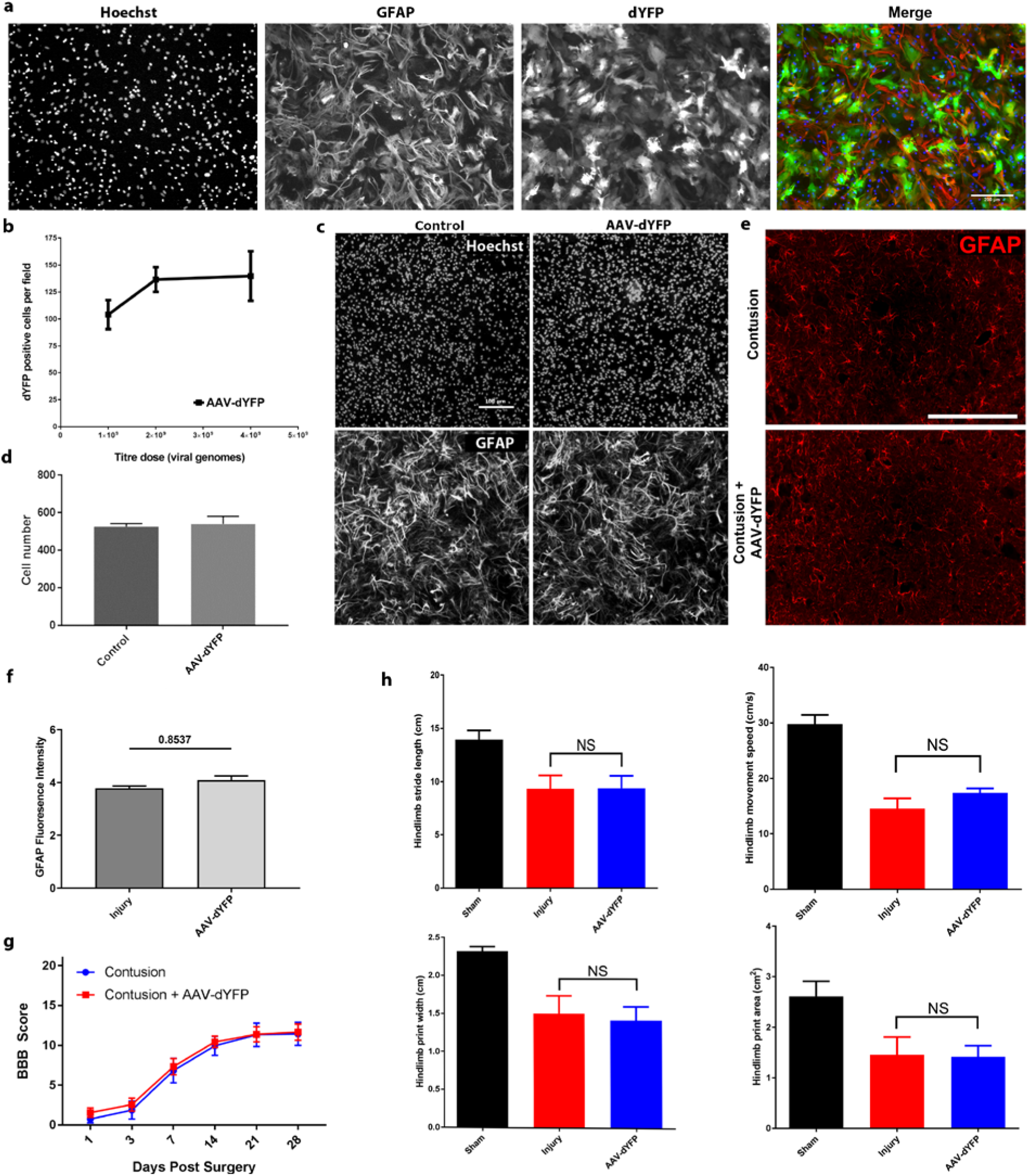
AAV-dYFP results in robust expression and does not affect astrogliosis or behavioural parameters. An astrocyte-selective AAV5 vector expressing dYFP under the expression of the GfaABC_1_D promoter (AAV-dYFP) resulted in robust transduction of primary spinal cord astrocytes (a, b; n = 4 independent cultures). Transduction did not affect astrogliosis or cell viability (c, d). In an *in vivo* spinal cord contusion model, AAV-dYFP did not affect GFAP immunostaining (scale bar = 200 μm); e, f), BBB locomotor scores (g) or CatWalk Gait XT analysis parameters (h; Sham n = 4, n = 9 for injury and AAV-dYFP groups).

Following *in vitro* experiments, we determined whether injections of the AAV-dYFP vector had any effects on astrogliosis or caused behavioural deficits in a spinal cord contusion model. This study was carried out over 4 weeks because beyond this time point any effect to behaviour or astrogliosis could reflect dYFP transgene-related effects rather than a response to the viral vector particles (Vandamme et al., 2017). As observed *in vitro*, AAV-dYFP did not affect astrogliosis in comparison to non-injected animals (Fig. 2 e, f). We further investigated whether the AAV-dYFP vector influenced behavioural measurements. As we observed previously (Griffin et al., 2019), AAV-dYFP intraspinal injections did not affect BBB locomotor score, or several parameters measured by CatWalk Gait XT analysis (Fig. 2 g, h).

### AAV-ADAMTS4 transduces primary spinal cord astrocytes and degrades TGFβ1-induced CSPGs

We conducted two proof-of-concept experiments to show that CSPGs are a substrate for ADAMTS4-mediated proteolysis (Fig. S3) and that ADAMTS4 can reverse the CSPG-induced inhibition of neurite number and length independent of cell viability (Fig. S4). Following packaging of AAV-ADAMTS4 we first tested whether the vector was functional in primary spinal cord astrocyte cell culture. TGFβ1 has previously been shown to increase CSPG expression in primary astrocyte culture (Smith and Strunz, 2005) and we investigated whether AAV-ADAMTS4 degrades TGFβ1-induced increases in CSPG content. Immunocytochemistry was used to visualise the expression of all GAGs and CS-GAGs specifically (WFA and CS56 staining, respectively). After exposure to TGFβ1, astrocytes produced large amounts of CS-GAG, and all GAGs (*P* < 0.0001; Fig. 3). Astrocytic expression of ADAMTS4 was low in the control condition and TGFβ1 treated cells (Fig. 3 a, b). Transduction with 4×10^9^ or 8×10^9^ vg AAV-ADAMTS4 resulted in a significant, three-fold increase in the expression of ADAMTS4 (*P* < 0.05). Equal ADAMTS4 expression was achieved by either 4×10^9^ or 8×10^9^ vg, indicating saturation of the cultures similar to what is observed by AAV-dYFP expression in cultures. Degradation of TGFβ1-induced-CSPGs by AAV-ADAMTS4 was visually evident. Both 4×10^9^ and 8×10^9^ vg AAV-ADAMTS4 reduced the fluorescence values for CS-GAGs and GAGs (*P* < 0.001; Fig. 3 c, d). GAG content was reduced to levels comparable to control whilst CS-GAG expression was reduced to slightly more than half that observed by TGFβ1 alone. 8×10^9^ vg degraded slightly more CS-GAG and GAG content compared to 4×10^9^ vg, though these differences were not significant. Recombinant ADAMTS4 enzyme (20 nM) was used as a positive control and significantly reduced levels of CS-GAG and GAG (*P* < 0.0001).

**Figure 3:**
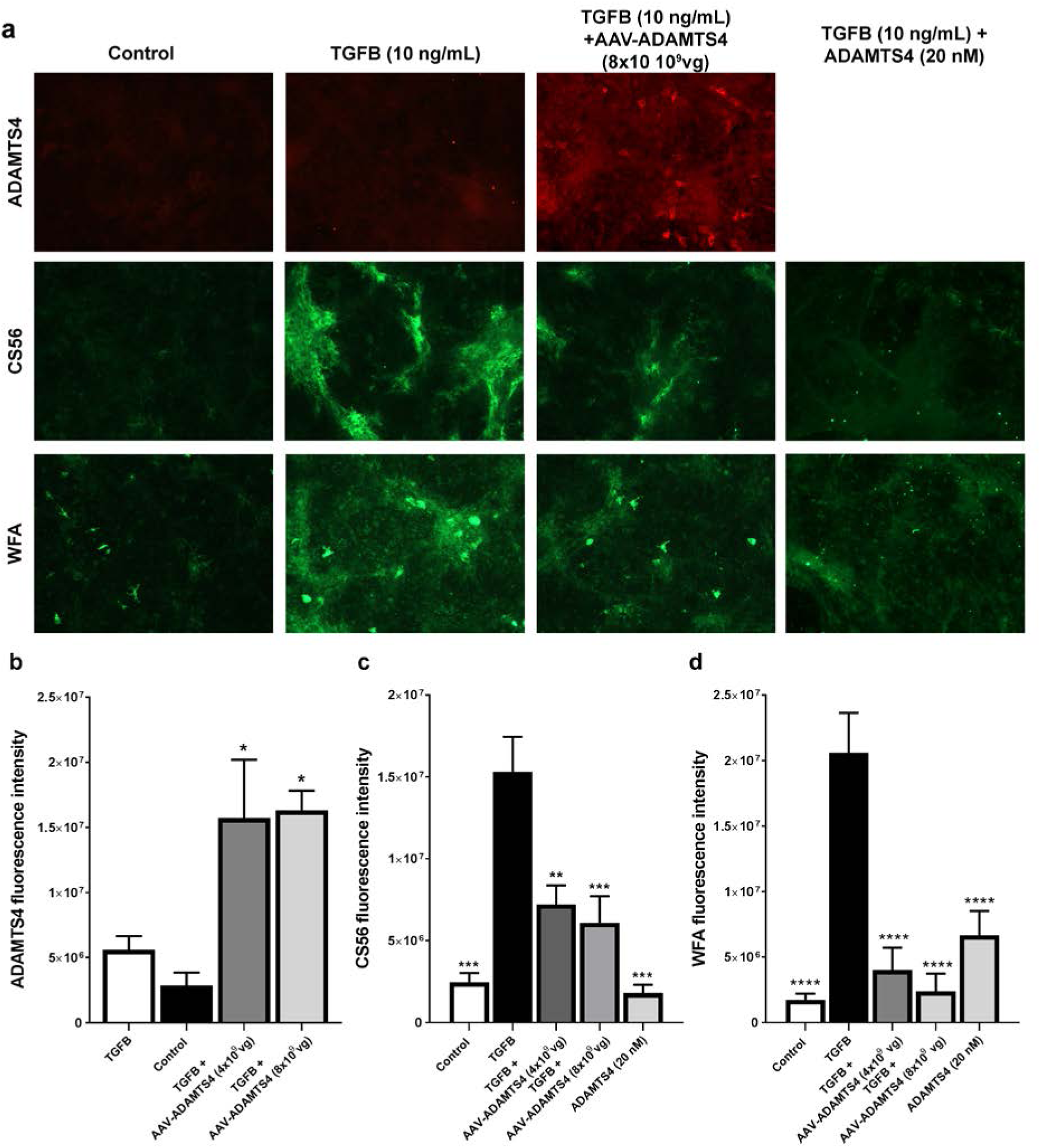
AAV-ADAMTS4 transduces primary spinal cord astrocytes and degrades CSPG components. Primary spinal cord astrocytes were transduced with the AAV-ADAMTS4 vector (either 4×10^9^ or 8×10^9^ vg) and stimulated with 10 ng/mL TGFβ1 two days later. The cells were fixed three days after stimulation and fluorescent immunocytochemistry was then used to detect the ADAMTS4 transgene (ADAMTS4), chondroitin sulphate (CS56) and glycosaminoglycan (WFA). a) 10x magnification images were captured using a Nikon TE-2000-U microscope. Images were thresholded and arbitrary fluorescence units were measured for b) ADAMTS4, c) CS56 and d) WFA. Data represents mean ± SEM (n = 4 independent cultures). A one-way ANOVA followed by a Tukey’s multiple comparisons test was used to test statistically significant differences *P < 0.05, **P < 0.001, ***P < 0.0001.

### ADAMTS4 transgene is expressed and functional in the rat spinal cord after infusion of AAV-ADAMTS4

We next investigated whether the ADAMTS4 transgene could be expressed *in vivo* in a contusion model of SCI. Immunohistochemistry was used to detect the ADAMTS4 transgene expression in the thoracic spinal cord ten weeks after thoracic SCI and infusions of AAV-ADAMTS4. Efficient ADAMTS4 expression was detected in the thoracic cord in sections 600-1200 µm rostral of the lesion edge compared to untransduced spinal cord (Fig. 4 a). Confocal microscopy was used to visualise ADAMTS4 expression and colocalisation with GFAP (Fig. 4 b). Basal ADAMTS4 expression was present in spinal cords that were not injected with the vector but this expression is at low levels. Expression was largely diffuse in grey and white matter, although staining is present around the cell bodies of neurons. We observed what appeared to be increased GFAP expression in spinal cords injected with AAV-ADAMTS4 compared to injury-only control (Fig. 4 b), To determine the cause of this we compared GFAP expression in control cords and those injected with AAV-dYFP and AAV-ADAMTS4 for 4 weeks, which indicated that ADAMST4 itself may induce astrogliosis that is not related to the viral vector or the injections (P < 0.05; Fig 4 b-d).

**Figure 4:**
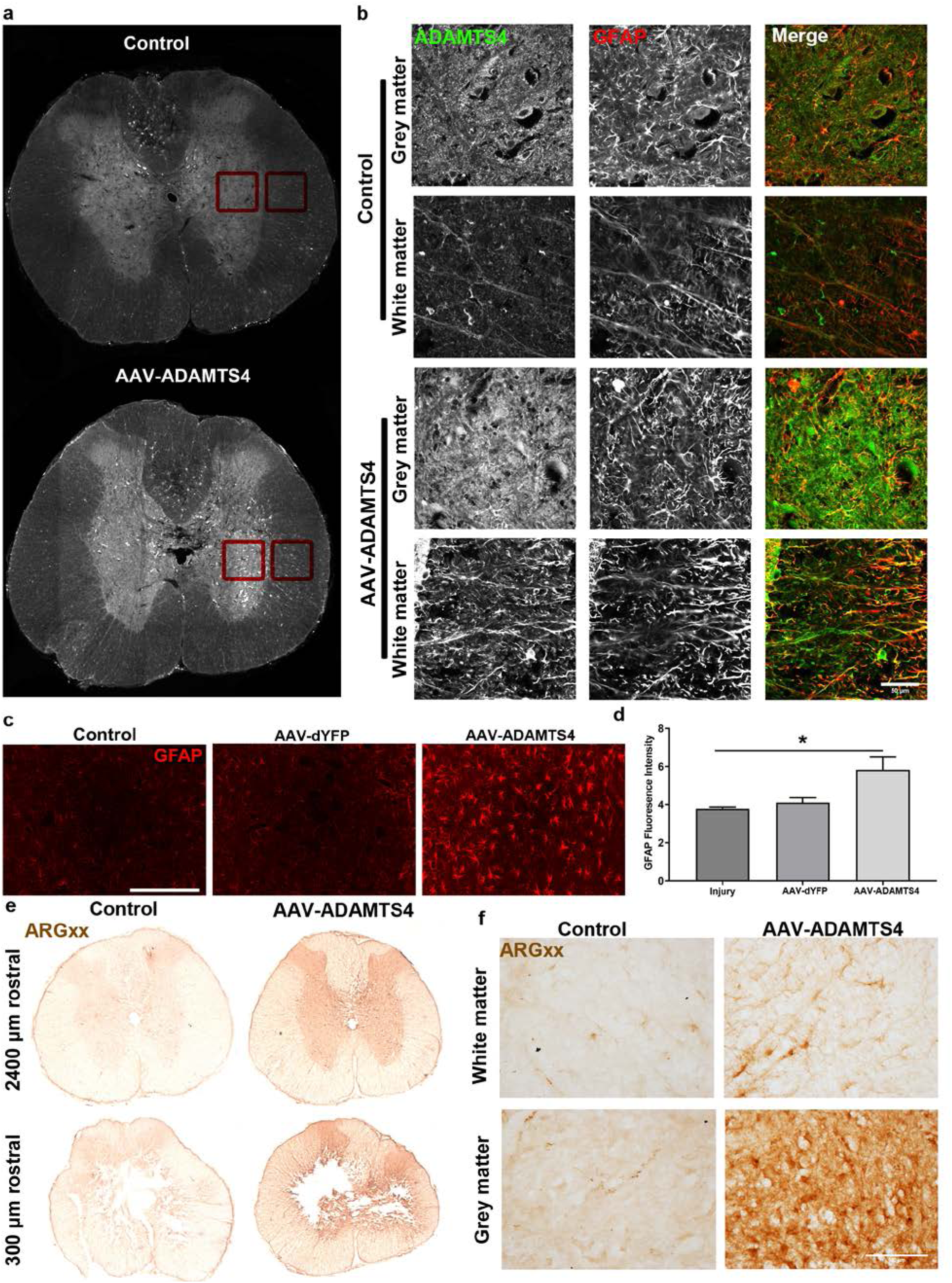
Intraspinal AAV-ADAMTS4 injections increase ADAMTS4 expression and ADAMTS4-specific degradation of aggrecan. Spinal cords were processed for immunohistochemistry to detect GFAP (red) and ADAMTS4 (green) expression in sections neighbouring the lesion (approximately 600-1200 µm rostral of the lesion edge). An EVOS FL Auto was used to capture scans of whole sections (a). An Olympus FV1000 microscope was used to capture images at 60x magnification, using conserved capture settings, in grey and white matter regions (b). Images are representative images. Scale bar = 50 µm. Increased ADAMTS4 expression is evident in the spinal cord of animals infused with AAV-ADAMTS4. Injections of AAV-ADAMTS4 increased GFAP expression compared to AAV-dYFP or to injury-only controls (scale bar = 200 μm; c, d). DAB immunohistochemistry was used to detect the neoepitope (ARGxx) created by ADAMTS-specific-proteolysis of aggrecan. A Leica DMR microscope was used to capture images with conserved capture settings at 2.5x (e). Large increases in the neoepitope were present in of spinal cords injected with the vector compared to un-injected cords. Images are representative images at either the lesion center or 2400 µm rostral to the edge of the lesion. Images at 25x magnification were captured in the grey matter of untreated and treated animals (f). Images are representative images. Scale bar = 100 µm.

We next determined if the ADAMTS4 transgene was functional in transduced spinal cord cells. An antibody against an ADAMTS-specific neoepitope (ARGxx) was used to serve as a proxy for increased ADAMTS4 expression and functional activity. DAB immunohistochemistry revealed robust ADAMTS4-aggrecanase activity throughout large volumes of spinal cord tissue (more than 4800 µm) following injection of AAV-ADAMTS4 (Fig. 4 e, f Fig S5). White matter ARGxx staining appeared in morphology to be localised to astrocytes. Staining was more intense in grey matter, particularly around the cell bodies of neurons (Fig. 4 f), which is consistent with the expression of aggrecan (Galtrey et al., 2008). Only faint or no staining was evident in untransduced tissue.

### AAV-ADAMTS4 treatment reduces lesion size after spinal cord injury

Luxol fast blue and eosin staining was used for stereological quantification of the lesion size and areas of grey and white matter following AAV injection with and without rehabilitation (Fig. 5). Force measurements from the contusion impaction device confirmed injuries were equal in severity across treatment groups. In AAV-ADAMTS4 transduced animals, a significant decrease in lesion size, increased white matter area, and increased total tissue area compared to contusion injury alone was observed (Fig. 5 e-g). Unexpectedly, tissue from animals that received rehabilitation alone appeared to have a larger lesion size compared to the unrehabilitated, transduced and untransduced animals (Fig. 5 a-e). The size of the whole sections from the rehabilitation groups (excluding the lesion area) is not significantly larger than the total tissue area of sham animals from the corresponding section. Instead, tissue from rehabilitated animals had a more rounded structure whereas the lesion center of unrehabilitated animals appears to be collapsed inwards (Fig. 5 a-d). Therefore, exercise rehabilitation may influence the shape and structure of the spinal cord, rather than affect lesion or tissue area.

**Figure 5:**
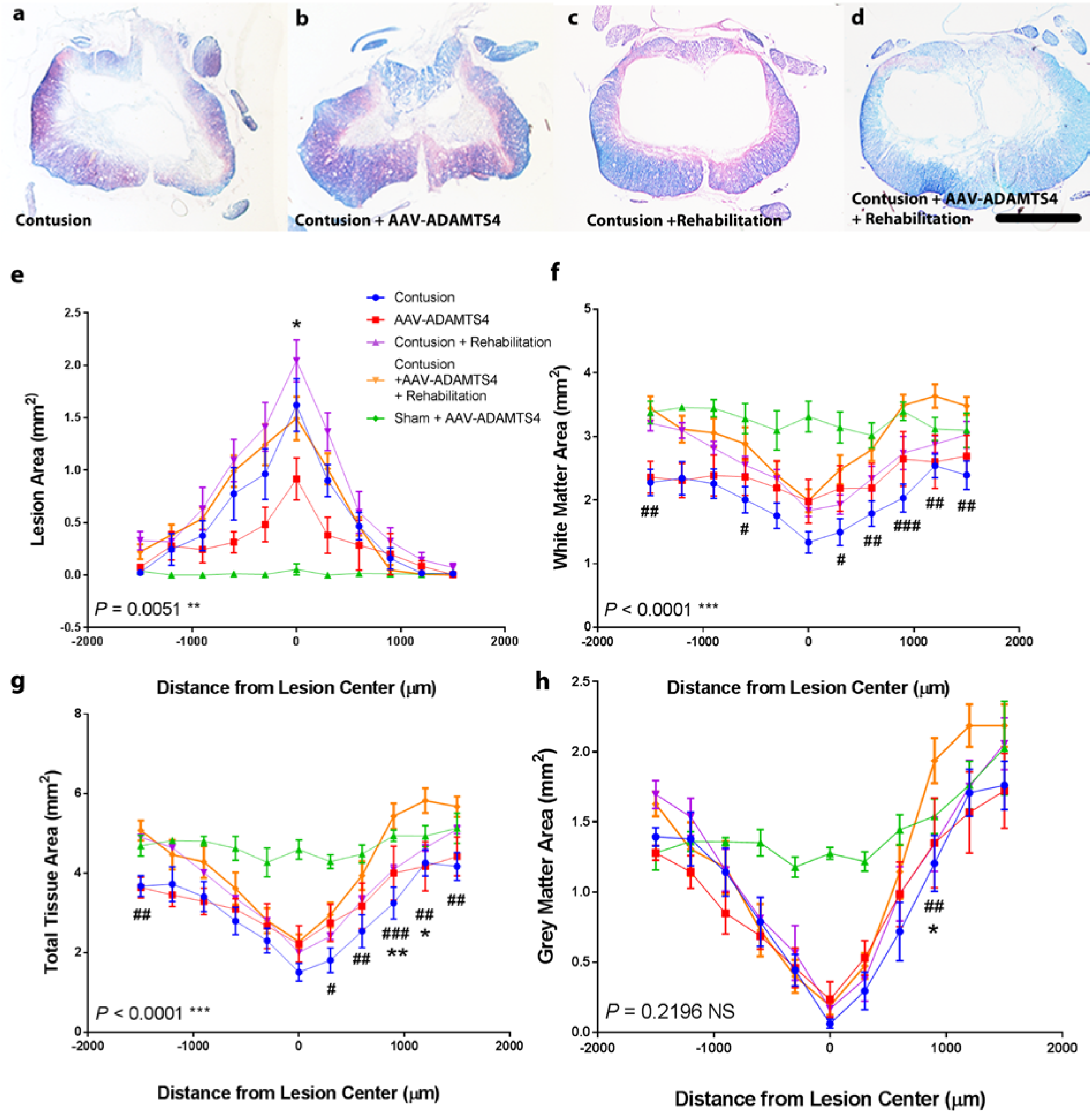
ADAMTS4 gene therapy leads to significantly reduced lesion size. Luxol fast blue-eosin histology was used to investigate the integrity of spinal cord tissue following contusion (a), contusion and AAV-ADAMTS4 treatment (b), contusion with rehabilitation (c), and contusion with AAV-ADAMTS4 treatment and rehabilitation (D). Staining was used to determine the area of the lesion (e), white matter (f), total remaining tissue (g) and grey matter (h). Data represent mean ± SEM (contusion n = 10, contusion + AAV-ADAMTS4 n = 9; contusion + rehabilitation n = 12; contusion + AAV - ADAMTS4 + rehabilitation n = 9, sham + AAV-ADAMTS4 n = 4; scale = 1 mm). A Two-Way ANOVA and Tukey’s multiple comparisons test were used to determine statistical significances where # compares contusion to contusion plus AAV-ADAMTS4 plus rehabilitation; *compares contusion plus rehabilitation to contusion plus AAV-ADAMTS4 plus rehabilitation. Results from the ANOVA tests comparing the variable of ‘treatment’ between the contusion plus rehabilitation to contusion plus AAV-ADAMTS4 plus rehabilitation are displayed on the figures. For all figures: NS = Non significant, *^/#^*P* < 0.05, **^/##^*P* < 0.001, ***^/###^*P* < 0.0001

### AAV-ADAMTS4 stimulates collateral sprouting of CST axons in the damaged spinal cord and increases serotonergic fiber density caudal to the lesion

A known consequence of CSPG digestion in the spinal cord is enhanced axonal sprouting and regeneration (Starkey et al., 2012). Therefore we investigated whether AAV-ADAMTS4 could enhance sprouting of hindlimb CST axons rostral to the lesion site through anterograde tracing. Intense BDA staining within neuron cell bodies and their axons was confirmed in the hindlimb motor cortex spanning -0.5 mm to -2.0 mm posterior in reference to Bregma (the coordinates for hindlimb neurons; data not shown (Neafsey et al., 1986). Hindlimb CST axons were detected in the thoracic spinal cord, 600-1200 µm rostral to the lesion edge. Axons leaving the dorsal CST bundle and entering grey matter territories were observed in both treated and untreated animals but more sprouting was observed in animals that received AAV-ADAMTS4 (red arrows; Fig 6 a). To quantify the level of axonal sprouting in all treatment groups, the total length of BDA labelled collateral sprouts outside of the CST were traced and measured (Fig. 6 b). In untransduced animals, loaded axons were mainly confined to the CST, whereas for animals that received AAV-ADAMTS4, axon sprouting and growth widespread through grey matter. The average sum of the length of axons traced for untransduced animals was 4369.83 µm ± 937.67 µm. This is comparable to untransduced animals that also received rehabilitation (4098.25 µm ± 2398.46 µm). The average sum of the length of axons traced for animals treated with AAV-ADAMTS4 (12,523.94 µm ± 3303.38 µm) was significantly different from animals that received the contusion alone (*P* < 0.0002). Animals that received AAV-ADAMTS4 + rehabilitation showed a significant increase in the sum of traced axons compared to both transduced groups (9576.31 µm ± 4160.09 µm; *P* < 0.05).

**Figure 6:**
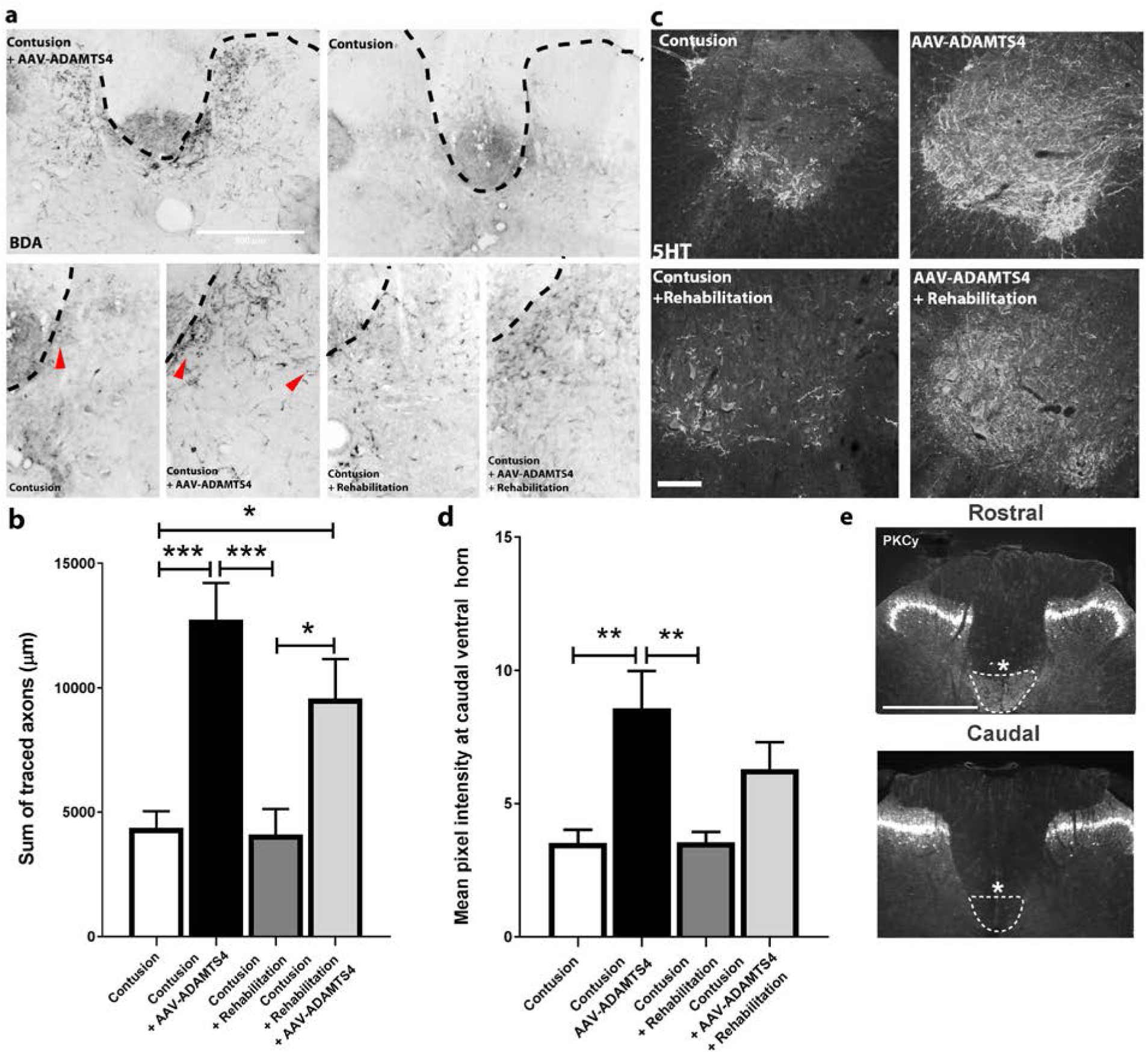
AAV-ADAMTS4 increases hindlimb CST collateral sprouting rostral to the lesion and serotonergic innervation caudal to the lesion. BDA was injected into the hindlimb motor cortex of rats 10 weeks after SCI. a) In untransduced animals, loaded axons were mainly confirmed to the CST 600-1200 µm rostral to the lesion edge, whereas treated animals displayed axon sprouting and growth widespread through grey matter (examples indicated by the red arrows; scale bar = 500 µm). ImageJ was used to quantify the total length of traced axons outside of the CST; transduced animals had a significantly greater amount of traced axons outside of the CST. b) Data represents the mean ± SEM. A two-tailed unpaired t-test was used to determine statistical differences between the two groups where \**P* < 0.05 (n = 7 for contusion + rehabilitation and contusion + AAV-ADAMTS4 + rehabilitation; n = 4 for contusion and contusion + AAV-ADAMTS4). c) Serotonergic fibers in the ventral horns of sections caudal to the injury (three sections: 600, 1200 and 1800 µm caudal to the injury were used; scale bar = 200 µm) were visualised. d) Images were consistently thresholded and the mean pixel intensity of serotonin immunoreactivity at the ventral horn was measured. Data represent the mean ± SEM (contusion n = 10; contusion + AAV-ADAMTS4 n = 7; contusion + rehabilitation n = 10; contusion + rehabilitation + AAV-ADAMTS4 n = 6). A one-way ANOVA followed by a Tukey’s multiple comparisons test was used to test statistically significant differences where \**P* < 0.05, \*\**P* < 0.001. e) Protein kinase C gamma immunofluorescence confirmed that abolishment of the CST rostral to the lesion (indicated by *).

Serotonin (5HT) projections that descend from the raphe nuclei and terminate in the ventral horn of the spinal cord provide excitatory input to motor neurons. Loss of these projections correlates with locomotor dysfunction (Schmidt and Jordan, 2000). ChABC promotes sprouting of serotonergic fibers caudal to a SCI (Barritt et al., 2006) and so it is possible the ADAMTS4 could also. To investigate serotonergic fibers caudal to the injury, 5HT immunoreactivity was examined in the ventral horns. Dense 5HT fiber staining was apparent in the ventral horns of AAV-ADAMTS4 transduced animals (Fig. 6 c, d) In contrast, significantly less 5HT immunoreactivity was observed in the ventral horns of untransduced animals as well as untransduced animals that received rehabilitation (*P* < 0.001 for both). The mean pixel intensity values of 5HT staining in both untransduced groups were nearly identical (3.52 ± 1.51 vs. 3.54 ± 1.21), indicating that rehabilitation did not affect these serotonergic tracts in the absence of AAV-ADAMTS4 expression.

We used a thoracic contusion SCI model that ablates the dorsal CST. The motor cortex and CST plays a critical role in manual dexterity of skilled behavioural tasks (Whishaw et al., 1998), and CSPG degradation using ChABC leads to neuroplasticity of the CST that is associated with recovery of such skilled behaviour tasks (Bradbury et al., 2002). We assessed whether there was a complete loss of the dorsal CST in animals following lesioning and therefore improvements to the horizontal error ladder test (a more skilled behaviour test) may be associated with adaptive plasticity of the CST due to ADAMTS4 activity. Complete lesioning of the dorsal CST was verified by staining spinal cord section rostral and caudal to the injury for protein kinas C gamma (PKCγ; Fig. 6 e)

### AAV-ADAMTS4 combined with hindlimb rehabilitation enhances functional recovery after spinal cord injury

CSPG degradation through the application of ChABC has been shown in a variety of animal models to improve behavioural outputs following spinal cord injury. We therefore investigated whether AAV-ADAMTS4 gene therapy could result in a similar observation using the BBB scoring system and horizontal error ladder test. Force measurements from the contusion impaction device confirmed injuries were equal in severity across treatment groups (Fig. 7 a). As a measure of overall health status, the AAV-ADAMTS4 vector did not negatively affect animal bodyweight (Fig 7 b). Animals in the AAV-ADAMTS4 group displayed significantly higher mean body weights compared to the contusion-only group at days 63 and 70 (two-way ANOVA with Bonferroni’s multiple comparisons test; shown with *; *P* < 0.05; *P* < 0.001). Animals that received rehabilitation recovered body weight at a significantly slower rate than those that were not rehabilitated from day 27 onwards (*P* < 0.05; symbols not shown).

**Figure 7:**
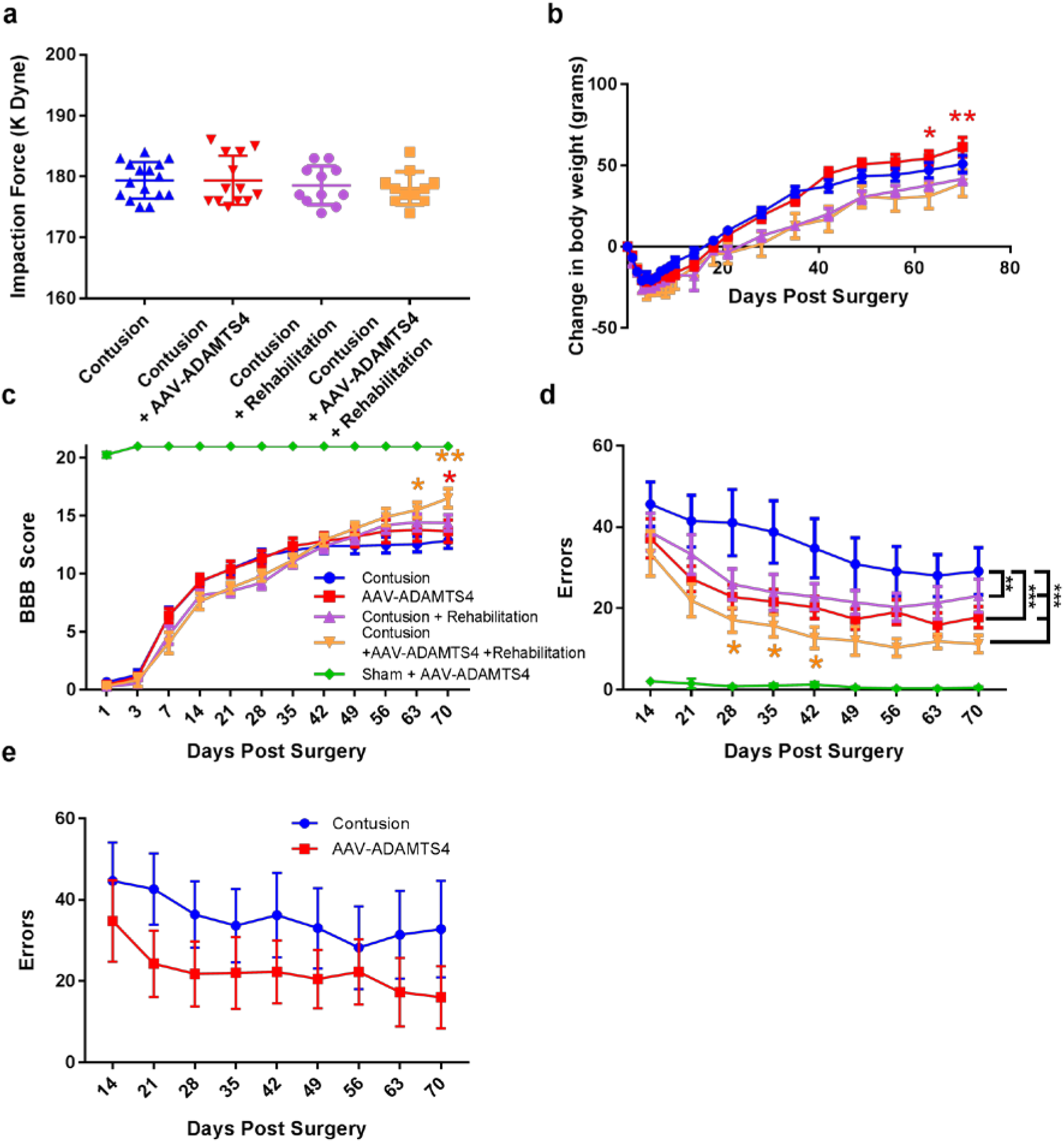
ADAMTS4 gene therapy with rehabilitation after thoracic spinal cord injury enhances improvements in behavioural assessments. The measured impaction force did not differ between treatment groups, confirming consistent injuries across all groups (a). The animals were weighed weekly as an indication of health status (b). BBB scores were measured for 10 weeks following the injury (c). Horizontal ladder testing began two weeks after injury where animals were recorded as they crossed a 1-meter horizontal ladder with irregular rungs. Data reported is the sum of the number of foot slips made for three runs. For (a) and (b), the data represents the mean ± SEM (contusion n = 14, contusion + AAV-ADAMTS4 n = 13; contusion + rehabilitation n = 12; contusion + AAV -ADAMTS4 + rehabilitation n = 9, sham + AAV-ADAMTS4 n = 4). For (a), a two-way, repeated measure ANOVA was used to determine significant differences between the groups over time and a Bonferroni multiple comparisons test was used to determine differences at each time point. For (b), since testing began two weeks after SCI and AAV injections, a non-repeated measure two-way ANOVA was used with Bonferroni multiple comparisons \**P* <0.05, \*\**P* <0.001, \*\*\**P* <0.0001: specifically. For (b), yellow * compares contusion + AAV -ADAMTS4 + rehabilitation with contusion and red * compares contusion + AAV -ADAMTS4 + rehabilitation to contusion + AAV -ADAMTS4. The results from AAV-ADAMTS4-treated animals compared to contusion-only animals was replicated in a repeat study of a smaller cohort showing that the result is replicable (e, n = 5 for both groups; data represent mean ± SEM; two-way ANOVA where \*\**P* <0.001).

For BBB scores, all of the animals displayed a BBB score of less than two (2 = slight movement of one or two joints) one day after spinal cord contusion confirming a consistent injury for all animals (Fig. 7 c). Though not statistically significant, between days 14-35 after injury, both rehabilitation groups showed lower BBB scores than their un-rehabilitated counterparts. From 35 days onwards, however, both rehabilitated groups displayed higher mean BBB values compared to the contusion only group (without rehabilitation). Animals that received either AAV-ADAMTS4 or rehabilitation alone showed higher BBB scores compared to the contusion-only group, although this is not a significant difference. Animals that received AAV-ADAMTS4 in combination with hindlimb rehabilitation showed significantly greater BBB scores compared to the contusion only group at days 63 and 70 (15.5 ± 1.9 vs. 13.0 ± 3.6, *P* < 0.05; 16.5 ± 2.4 vs. 13.2 ± 3.8, *P* < 0.001). Interestingly, at days 63 and 70, those in the AAV-ADAMTS4 group displayed significantly greater body weights than their contusion-only counterparts (Fig 7 b). Furthermore, animals that received AAV-ADAMTS4 in combination with hindlimb rehabilitation showed significantly greater BBB scores at day 70 compared to animals that received AAV-ADAMTS4 without rehabilitation (16.5 ± 2.4 vs. 14.4 ± 2.4, *P* < 0.05).

Functional effects were also assessed using the horizontal ladder test, a skilled motor task. Two weeks after SCI and vector injections, a large number of errors were made by all groups. Animals that received contusion alone had more errors compared to those treated with the vector (45.60 ± 21.15 vs. 37.23 ± 17.36; Fig. 7 d) which may indicate that the therapeutic benefit of the ADAMTS4 vectors appears earlier than two weeks. For all groups, fewer errors were made over the course of the following five weeks at which point the number of errors appears to plateau for the remaining three weeks of scoring. At the end of the 10 weeks, rats that received AAV-ADAMTS4 with rehabilitation were making less than half the number of errors made by rats without treatment or rehabilitation (11.22 ± 6.4 vs 29.13 ± 22.41). Over the course of testing, the addition of AAV-ADAMTS4 alone or in combination with rehabilitation significantly improved function compared to contusion only (*P* < 0.0001; two-way ANOVA). Similarly, rehabilitation alone led to significantly improved function (*P* < 0.001; two-way ANOVA).

Replication of experimental results is increasingly being considered crucial for translation of research findings. In order to confirm the effect of the ADAMTS-4 gene therapy a behavioural study with a smaller cohort was carried out using the AAV-ADAMTS and contusion-only groups. Even with a smaller cohort an effect of the gene therapy on behavioural improvement was seen, confirming our results and the potential of AAV-ADAMTS4 gene delivery (Fig 7 e).

## Discussion

Here we demonstrate that an AAV vector expressing human ADAMTS4 under the control of an astrocytic promoter is capable of transducing spinal cord astrocytes and degrading CSPGs. This limited lesion size, promoted sprouting of the hindlimb CST, increased caudal serotonergic fiber density, and lead to an improvement in hindlimb locomotor ability in a contusive experimental model that closely mimics human traumatic SCI. We further showed that hindlimb-specific rehabilitation leads to an enhanced improvement in functional recovery.

AAV-ADAMST4 transduction of cultured primary astrocytes resulted in a large increase in ADAMTS4 expression compared to untransduced astrocytes, or astrocytes treated with TGFβ1. Significant decreases in both chondroitin sulphate and glycosaminoglycan content was present in transduced cells. Following the confirmation that the AAV-ADAMTS4 vector is functional in cultures, it was then delivered into the rat spinal cord in a contusion injury model to investigate whether it could be therapeutic. AAV-ADAMTS4 enabled efficient expression of the enzyme for at least 13 weeks. ADAMTS’s are expressed throughout the CNS, and ADAMTS4 appears to be the most highly expressed in the CNS under basal conditions (Yuan et al., 2002) - although, literature pertaining to the regional and cellular expression, as well as the biological functions of ADAMTS4 in the CNS is limited. ADAMTS4 has been found to be expressed in spinal cord astrocytes and microglia, as well as neurons, however, the relative levels of ECM expression were not described (Cross et al., 2006; Yuan et al., 2002). In untreated injured tissue, we observed low, diffuse immunostaining of ADAMTS4 not localised to any specific cell population. This is not unexpected as it is known to be released from cells to act on the ECM (Gottschall and Howell, 2015). The large increase in ADAMTS4 expression with transduction resulted in cellular staining as well as an increased expression within the ECM. Expression was present in astrocytes but neuronal expression was also evident. Potentially this could reflect localisation to CSPG-rich surroundings of neuronal cell bodies. In this study, an antibody was used that recognises an ADAMTS4 N-terminal neoepitope (ARGxx) produced after cleavage between amino acids EGE and ARG within the interglobular domain of aggrecan. AAV-ADAMTS4 led to a dramatic increase in this neoepitope expression and was widespread throughout the spinal cord. Aggrecan expression, however, is known to be mainly localised to the grey matter in adult spinal cord tissue where expression is present in the ECM and found on all PNN-bearing neurons (Galtrey et al., 2008). Consistent with the present study, staining was greater in grey matter as opposed to white matter and the localisation of staining appeared to be present in the ECM and localised to the cell bodies of neurons in the grey matter.

AAV-ADAMTS4 increased GFAP expression *in vivo* that was not related to the injection or to the viral vector as no such increased was observed after injection of AAV-dYFP. Reports of ADAMTS4 affecting astrogliosis or neuroinflammation are limited to two studies. ADAMTS4 has been reported to have anti-inflammatory effects in a mouse model of ischemic stroke in terms of reducing a variety of pro-inflammatory cytokines as well as reducing astrogliosis and macrophage infiltration (Lemarchant et al., 2016a). Conversely, recombinant ADAMTS4 increased astrogliosis in the lumbar spinal cord female SOD1 mice which was associated with neurodegeneration in a mouse model of amyotrophic later sclerosis (Lemarchant et al., 2016b). Astrogliosis and astrocytic scars are widely regarded to be casual in the failure of mature CNS axon regeneration, although, our understanding of the role of astrogliosis in regeneration is changing. This has become evident through a landmark paper which proved that activated, scar-forming astrocytes aids CNS axon regeneration and preventing astrocytic scar formation significantly reduced stimulated axon growth (Anderson et al., 2016). In this study, RNA sequencing revealed that activated astrocytes in SCI lesion express multiple axon-growth supporting molecules. Therefore, ADAMTS4 may have beneficial effects through promoting astrogliosis but further investigation is required to tease out the therapeutic mechanism of AAV-ADAMTS4.

Compensatory spinal networks form spontaneously after SCI and correlate with functional recovery in motor systems (Bareyre et al., 2004; Rosenzweig et al., 2010; Weidner et al., 2001). These plasticity events are enhanced through the application of anti-inhibitory treatments such as ChABC or anti-NogoA antibody treatment by removing inhibitory factors in the ECM (Barritt et al., 2006; Gonzenbach et al., 2012). It was therefore probable that AAV-ADAMTS4-mediated degradation of CPSGs is causing a similar effect. Sprouting of the CST axons descending from the hindlimb motor cortex was confirmed using BDA anterograde tracing whereby a significant increase in the number of traced-CST axons sprouting from the CST rostral to the lesion in animals that received AAV-ADAMTS4 injections was seen. Many similar observations have been made after the removal of inhibitory myelin components or CSPGs (Barritt et al., 2006; Bradbury et al., 2002; Garcia-Alias et al., 2009; Li et al., 2004). AAV-ADAMTS4 treatment also led to significant plasticity of the intact descending raphe-spinal serotonergic system after SCI whereby a significant increase in serotonergic fiber density in the ventral horn caudal to the injury was witnessed. Spinal serotonin levels are known to facilitate locomotor function and may influence functional recovery after SCI (Schmidt and Jordan, 2000). Furthermore, increased innervation of the ventral horn by sprouting serotoninergic fibres after ChABC, LV-ChABC and anti-NogoA treatments have been associated with a recovery in locomotor function (Barritt et al., 2006; Bartus et al., 2014; Bregman et al., 1995; Li et al., 2004). An important consideration is that AAV-ADAMTS4 may have contributed to the serotonergic fiber increase through neuroprotective tissue preservation at the lesion epicentre. Whether by axonal sparing, or fiber sprouting, AAV-ADAMTS can increase serotoninergic fibers caudal to the injury. In this present study, we did not observe rehabilitation-induced CST or 5HT plasticity. This result is similar to several previous observations in which exercise training did not increase CST collateral sprouting or 5HT fiber density caudal to the lesion (Alluin et al., 2014). In another study, neither skilled forepaw rehabilitation or general rehabilitation in rats that received a C4 dorsal cut increased CST axon midline crossing or sprouting (Garcia-Alias et al., 2009). In fact, the combination of ChABC with rehabilitation led to less collateral crossing of BDA-loaded CST axons. Likewise, intense wheel-rehabilitation in mice following thoracic hemisection did not increase caudal serotonin immunofluorescence at 10 weeks post-SCI (Loy et al., 2018). This is complicated by the fact the contrary is also reported to occur (Engesser-Cesar et al., 2007; Maier et al., 2008). These differences may reflect the type of exercise prescribed. Ultimately, we have much more to learn how activity-influenced regeneration and adaptive plasticity.

Improvements in motor function have previously been reported after large-scale CSPG degradation mediated by a lentiviral ChABC gene therapy (Bartus et al., 2014). Similarly, we observed a significant functional recovery on the horizontal ladder following AAV-ADAMTS4 treatment without rehabilitation. This improvement occurred early and continued throughout the testing period. It is likely due to neuroprotective effects since these animals did not show the same level of impairment in the task at week two, despite having the same severity of injury. A similar observation was made for the lentiviral ChABC therapy whereby significant benefit (in terms of performance on error ladder task) was reported prior to one week after injury and vector injection (Bartus et al., 2014). The authors confirmed that LV-ChABC could reduce the progression of secondary injury pathology in early time points leading to decreased lesion size. LV-ChABC shifted macrophage polarisation to an M2 phenotype indicated by increased expression of CD206 that weren’t always associated with IBA1 staining (Bartus et al., 2014). Enhanced recovery is associated with infiltrating M2 macrophages leading to enhanced tissue recovery and repair following SCI (Shechter et al., 2013), which is hypothesised to be the reason that LV-ChABC results in decreased lesion size (Bartus et al., 2014). It was also shown in a mouse model of ischemic stroke, ADAMTS4 displayed anti-inflammatory and neuroregenerative roles (Lemarchant et al., 2016a). This mechanism may also explain how we observed reduced lesion size in animals that received AAV-ADAMTS4 therapy compared to those that did not.

AAV-ADAMTS4 treatment alone resulted in a significant improvement in the horizontal ladder test, however, this improvement could be considered modest and arguably not reflective of the changes seen histologically. The same has previously been observed after ChABC treatment in the spinal cord and it was hypothesised by Fawcett and colleagues that many of the new connections may be non-useful and that through using appropriate rehabilitative strategies could facilitate the strengthening of beneficial connections (Garcia-Alias et al., 2009). Therefore, hindlimb rehabilitation was included with AAV-ADAMTS4 treatment to determine if this would give an improved functional outcome. Indeed, for the BBB and horizontal ladder test, rehabilitated animals had significantly better motor function compared to animals that did not receive hindlimb rehabilitation. Furthermore, this was significantly enhanced with AAV-ADAMTS4 combined therapy. Between days 14-35 after SCI (one week after the onset of their intense rehabilitation program), rehabilitation appeared to worsen BBB scores compared to animals that weren’t subjected to rehabilitation (though this deficit was not observed in the horizontal ladder test). Interestingly, there are no other reports of treadmill training worsening BBB scores in rats after SCI. On the contrary, no improvements or early improvements to BBB scores following treadmill training have been reported by multiple teams (Fouad et al., 2000; Hayashibe et al., 2016; Heng and de Leon, 2009; Multon et al., 2003). Specifically, quadrupedal training did not affect BBB scores above ‘self-training’ (Fouad et al., 2000), whereas bipedal, hindlimb treadmill training was shown to improve BBB scores, in some cases with an improvement of more than eight points (Hayashibe et al., 2016; Multon et al., 2003). The deficits in BBB scores in the present study could be the result of specifically training hindlimb, bipedal locomotion. This makes intuitive sense since the BBB scoring system requires the assessment of quadrupedal parameters and training of only the hindlimbs could impact on the animal’s ability to regain quadrupedal coordination. By day 14 following a 175 KDyne contusion, animals would typically have regained at least occasionally coordinated movements. However, animals that received hindlimb rehabilitation appeared to be hindered at this milestone yet having greater paw function compared to control that allow them to perform better on the horizontal ladder test. After day 35, natural quadrupedal, self-training movements within their home cages may have allowed the animals to eventually recover inter-limb coordination. Following this achievement, animals that received both treatment and rehabilitation appear to have a further, linear improvement representing their previously masked improvements. Whereas, for rehabilitated animals without treatment BBB scores begin to plateau. The ability of specific rehabilitation regimes to improve on behaviour while negatively affecting another has been previously reported. For example, general rehabilitation in the form of an environmental enrichment cage was shown to negatively affect skilled tasks in rats following dorsal column injury (Garcia-Alias et al., 2009). Likewise, spinally transected cats could be trained in weight support or step, however, training one behaviour would extinguish the other (De Leon et al., 1998a, b). It is likely that here, specifically training the animals in bipedal locomotion hinders them from reaching the accomplishment of forelimb-hindlimb inter-coordination, whilst not affecting their performance at the ladder test. In the present study, AAV-ADAMTS4 did not lead to any observable detrimental effects, although, we do not know whether longer-term CSPG digestion by AAV-ADAMTS4 could do so. Therefore, in the future it may be optimal to regulate ADAMTS4 gene expression, for example by using an AAV vector with an inducible promoter.

To conclude, a novel AAV vector was created for gene delivery to spinal cord astrocytes. This vector system delivering ADAMTS4 promotes functional recovery after spinal cord injury by decreasing lesion size and enhancing neuroplasticity. Combining AAV-ADAMTS4 gene therapy with specific rehabilitation can further enhance functional improvements. Due to the unparalleled safety profile of AAV vectors, and the safety associated with using a human gene, along with evidence that combination approaches that include rehabilitation are likely to give the best outcomes for clinical treatment, this therapy provides a promising candidate for clinical translation.

## Abbreviations

AAV: Adeno-associated virus
ADAMTS: A disintegrin and metalloproteinase with thrombospondin motifs
ChABC: Chondroitinase ABC
CS-GAG: Chondroitin sulphate glycosaminoglycan
CSPG: Chondroitin sulphate proteoglycan
ECM: Extracellular matrix
GAG: Glycosaminoglycan
LV: Lentiviral vector

## Acknowledgments

We would like to greatly thank the CatWalk Spinal Cord Injury Research Trust for funding this research and the Maurice Phyllis Paykel Trust for funding the purchase of the rodent treadmill. We would like the thank Denise Griffin for sewing the jackets for the rats.

## References

Alluin, O., Delivet-Mongrain, H., Gauthier, M.K., Fehlings, M.G., Rossignol, S., Karimi-Abdolrezaee, S., 2014. Examination of the combined effects of chondroitinase ABC, growth factors and locomotor training following compressive spinal cord injury on neuroanatomical plasticity and kinematics. PLoS One 9, e111072.

Anderson, M.A., Burda, J.E., Ren, Y., Ao, Y., O’Shea, T.M., Kawaguchi, R., Coppola, G., Khakh, B.S., Deming, T.J., Sofroniew, M.V., 2016. Astrocyte scar formation aids central nervous system axon regeneration. Nature 532, 195–200.

Bareyre, F.M., Kerschensteiner, M., Raineteau, O., Mettenleiter, T.C., Weinmann, O., Schwab, M.E., 2004. The injured spinal cord spontaneously forms a new intraspinal circuit in adult rats. Nat Neurosci 7, 269–277.

Barritt, A.W., Davies, M., Marchand, F., Hartley, R., Grist, J., Yip, P., McMahon, S.B., Bradbury, E.J., 2006. Chondroitinase ABC promotes sprouting of intact and injured spinal systems after spinal cord injury. J Neurosci 26, 10856–10867.

Bartus, K., James, N.D., Didangelos, A., Bosch, K.D., Verhaagen, J., Yanez-Munoz, R.J., Rogers, J.H., Schneider, B.L., Muir, E.M., Bradbury, E.J., 2014. Large-scale chondroitin sulfate proteoglycan digestion with chondroitinase gene therapy leads to reduced pathology and modulates macrophage phenotype following spinal cord contusion injury. J Neurosci 34, 4822–4836.

Basso, D.M., Beattie, M.S., Bresnahan, J.C., 1995. A sensitive and reliable locomotor rating scale for open field testing in rats. Journal of neurotrauma 12, 1–21.

Bradbury, E.J., Carter, L.M., 2011. Manipulating the glial scar: chondroitinase ABC as a therapy for spinal cord injury. Brain Res Bull 84, 306–316.

Bradbury, E.J., Moon, L.D., Popat, R.J., King, V.R., Bennett, G.S., Patel, P.N., Fawcett, J.W., McMahon, S.B., 2002. Chondroitinase ABC promotes functional recovery after spinal cord injury. Nature 416, 636–640.

Bregman, B.S., Kunkel-Bagden, E., Schnell, L., Dai, H.N., Gao, D., Schwab, M.E., 1995. Recovery from spinal cord injury mediated by antibodies to neurite growth inhibitors. Nature 378, 498–501.

Burnside, E.R., De Winter, F., Didangelos, A., James, N.D., Andreica, E.C., Layard-Horsfall, H., Muir, E.M., Verhaagen, J., Bradbury, E.J., 2018. Immune-evasive gene switch enables regulated delivery of chondroitinase after spinal cord injury. Brain.

Chen, K., Marsh, B.C., Cowan, M., Al’Joboori, Y.D., Gigout, S., Smith, C.C., Messenger, N., Gamper, N., Schwab, M.E., Ichiyama, R.M., 2017. Sequential therapy of anti-Nogo-A antibody treatment and treadmill training leads to cumulative improvements after spinal cord injury in rats. Exp Neurol 292, 135–144.

Cross, A.K., Haddock, G., Surr, J., Plumb, J., Bunning, R.A., Buttle, D.J., Woodroofe, M.N., 2006. Differential expression of ADAMTS-1, -4, -5 and TIMP-3 in rat spinal cord at different stages of acute experimental autoimmune encephalomyelitis. Journal of autoimmunity 26, 16-23.

Curcio, M., Bradke, F., 2018. Axon Regeneration in the Central Nervous System: Facing the Challenges from the Inside. Annu Rev Cell Dev Biol 34, 495–521.

De Leon, R.D., Hodgson, J.A., Roy, R.R., Edgerton, V.R., 1998a. Full weight-bearing hindlimb standing following stand training in the adult spinal cat. J Neurophysiol 80, 83–91.

de Leon, R.D., Hodgson, J.A., Roy, R.R., Edgerton, V.R., 1998b. Locomotor capacity attributable to step training versus spontaneous recovery after spinalization in adult cats. J Neurophysiol 79, 1329–1340.

During, M.J., Young, D., Baer, K., Lawlor, P., Klugmann, M., 2003. Development and optimization of adeno-associated virus vector transfer into the central nervous system. Methods in molecular medicine 76, 221–236.

Engesser-Cesar, C., Ichiyama, R.M., Nefas, A.L., Hill, M.A., Edgerton, V.R., Cotman, C.W., Anderson, A.J., 2007. Wheel running following spinal cord injury improves locomotor recovery and stimulates serotonergic fiber growth. Eur J Neurosci 25, 1931–1939.

Fawcett, J.W., Curt, A., 2009. Damage control in the nervous system: rehabilitation in a plastic environment. Nat Med 15, 735–736.

Fouad, K., Metz, G.A., Merkler, D., Dietz, V., Schwab, M.E., 2000. Treadmill training in incomplete spinal cord injured rats. Behavioural brain research 115, 107–113.

Galtrey, C.M., Kwok, J.C., Carulli, D., Rhodes, K.E., Fawcett, J.W., 2008. Distribution and synthesis of extracellular matrix proteoglycans, hyaluronan, link proteins and tenascin-R in the rat spinal cord. Eur J Neurosci 27, 1373–1390.

Garcia-Alias, G., Barkhuysen, S., Buckle, M., Fawcett, J.W., 2009. Chondroitinase ABC treatment opens a window of opportunity for task-specific rehabilitation. Nat Neurosci 12, 1145–1151.

Gonzenbach, R.R., Zoerner, B., Schnell, L., Weinmann, O., Mir, A.K., Schwab, M.E., 2012. Delayed anti-nogo-a antibody application after spinal cord injury shows progressive loss of responsiveness. Journal of neurotrauma 29, 567–578.

Gottschall, P.E., Howell, M.D., 2015. ADAMTS expression and function in central nervous system injury and disorders. Matrix Biol 44-46, 70-76.

Griffin, J.M., Fackelmeier, B., Fong, D.M., Mouravlev, A., Young, D., O’Carroll, S.J., 2019. Astrocyte-selective AAV gene therapy through the endogenous GFAP promoter results in robust transduction in the rat spinal cord following injury. Gene Ther.

Hayashibe, M., Homma, T., Fujimoto, K., Oi, T., Yagi, N., Kashihara, M., Nishikawa, N., Ishizumi, Y., Abe, S., Hashimoto, H., Kanekiyo, K., Imagita, H., Ide, C., Morioka, S., 2016. Locomotor improvement of spinal cord-injured rats through treadmill training by forced plantar placement of hind paws. Spinal cord 54, 521–529.

Heng, C., de Leon, R.D., 2009. Treadmill training enhances the recovery of normal stepping patterns in spinal cord contused rats. Experimental neurology 216, 139–147.

Kerstetter, A.E., Miller, R.H., 2012. Isolation and culture of spinal cord astrocytes. Methods in molecular biology 814, 93–104.

Lemarchant, S., Dunghana, H., Pomeshchik, Y., Leinonen, H., Kolosowska, N., Korhonen, P., Kanninen, K.M., Garcia-Berrocoso, T., Montaner, J., Malm, T., Koistinaho, J., 2016a. Anti-inflammatory effects of ADAMTS-4 in a mouse model of ischemic stroke. Glia 64, 1492–1507.

Lemarchant, S., Pomeshchik, Y., Kidin, I., Karkkainen, V., Valonen, P., Lehtonen, S., Goldsteins, G., Malm, T., Kanninen, K., Koistinaho, J., 2016b. ADAMTS-4 promotes neurodegeneration in a mouse model of amyotrophic lateral sclerosis. Mol Neurodegener 11, 10.

Li, S., Liu, B.P., Budel, S., Li, M., Ji, B., Walus, L., Li, W., Jirik, A., Rabacchi, S., Choi, E., Worley, D., Sah, D.W., Pepinsky, B., Lee, D., Relton, J., Strittmatter, S.M., 2004. Blockade of Nogo-66, myelin-associated glycoprotein, and oligodendrocyte myelin glycoprotein by soluble Nogo-66 receptor promotes axonal sprouting and recovery after spinal injury. J Neurosci 24, 10511–10520.

Loy, K., Bareyre, F.M., 2019. Rehabilitation following spinal cord injury: how animal models can help our understanding of exercise-induced neuroplasticity. Neural Regen Res 14, 405–412.

Loy, K., Schmalz, A., Hoche, T., Jacobi, A., Kreutzfeldt, M., Merkler, D., Bareyre, F.M., 2018. Enhanced Voluntary Exercise Improves Functional Recovery following Spinal Cord Injury by Impacting the Local Neuroglial Injury Response and Supporting the Rewiring of Supraspinal Circuits. J Neurotrauma 35, 2904–2915.

Maier, I.C., Baumann, K., Thallmair, M., Weinmann, O., Scholl, J., Schwab, M.E., 2008. Constraint-induced movement therapy in the adult rat after unilateral corticospinal tract injury. J Neurosci 28, 9386–9403.

Metz, G.A., Whishaw, I.Q., 2009. The ladder rung walking task: a scoring system and its practical application. J Vis Exp.

Mudannayake, J.M., Mouravlev, A., Fong, D.M., Young, D., 2016. Transcriptional activity of novel ALDH1L1 promoters in the rat brain following AAV vector-mediated gene transfer. Mol Ther Methods Clin Dev 3, 16075.

Muir, E.M., Fyfe, I., Gardiner, S., Li, L., Warren, P., Fawcett, J.W., Keynes, R.J., Rogers, J.H., 2010. Modification of N-glycosylation sites allows secretion of bacterial chondroitinase ABC from mammalian cells. Journal of biotechnology 145, 103–110.

Multon, S., Franzen, R., Poirrier, A.L., Scholtes, F., Schoenen, J., 2003. The effect of treadmill training on motor recovery after a partial spinal cord compression-injury in the adult rat. Journal of neurotrauma 20, 699–706.

Naso, M.F., Tomkowicz, B., Perry, W.L., 3rd, Strohl, W.R., 2017. Adeno-Associated Virus (AAV) as a Vector for Gene Therapy. BioDrugs 31, 317-334.

Neafsey, E.J., Bold, E.L., Haas, G., Hurley-Gius, K.M., Quirk, G., Sievert, C.F., Terreberry, R.R., 1986. The organization of the rat motor cortex: a microstimulation mapping study. Brain research 396, 77–96.

Rosenzweig, E.S., Courtine, G., Jindrich, D.L., Brock, J.H., Ferguson, A.R., Strand, S.C., Nout, Y.S., Roy, R.R., Miller, D.M., Beattie, M.S., Havton, L.A., Bresnahan, J.C., Edgerton, V.R., Tuszynski, M.H., 2010. Extensive spontaneous plasticity of corticospinal projections after primate spinal cord injury. Nature neuroscience 13, 1505–1510.

Rudge, J.S., Silver, J., 1990. Inhibition of neurite outgrowth on astroglial scars in vitro. The Journal of neuroscience: the official journal of the Society for Neuroscience 10, 3594–3603.

Schlimgen, R., Howard, J., Wooley, D., Thompson, M., Baden, L.R., Yang, O.O., Christiani, D.C., Mostoslavsky, G., Diamond, D.V., Duane, E.G., Byers, K., Winters, T., Gelfand, J.A., Fujimoto, G., Hudson, T.W., Vyas, J.M., 2016. Risks Associated With Lentiviral Vector Exposures and Prevention Strategies. J Occup Environ Med 58, 1159–1166.

Schmidt, B.J., Jordan, L.M., 2000. The role of serotonin in reflex modulation and locomotor rhythm production in the mammalian spinal cord. Brain research bulletin 53, 689–710.

Shechter, R., Miller, O., Yovel, G., Rosenzweig, N., London, A., Ruckh, J., Kim, K.W., Klein, E., Kalchenko, V., Bendel, P., Lira, S.A., Jung, S., Schwartz, M., 2013. Recruitment of beneficial M2 macrophages to injured spinal cord is orchestrated by remote brain choroid plexus. Immunity 38, 555–569.

Smith, G.M., Strunz, C., 2005. Growth factor and cytokine regulation of chondroitin sulfate proteoglycans by astrocytes. Glia 52, 209–218.

Snow, D.M., Brown, E.M., Letourneau, P.C., 1996. Growth cone behavior in the presence of soluble chondroitin sulfate proteoglycan (CSPG), compared to behavior on CSPG bound to laminin or fibronectin. International journal of developmental neuroscience: the official journal of the International Society for Developmental Neuroscience 14, 331–349.

Snow, D.M., Lemmon, V., Carrino, D.A., Caplan, A.I., Silver, J., 1990. Sulfated proteoglycans in astroglial barriers inhibit neurite outgrowth in vitro. Experimental neurology 109, 111–130.

Snow, D.M., Letourneau, P.C., 1992. Neurite outgrowth on a step gradient of chondroitin sulfate proteoglycan (CS-PG). Journal of neurobiology 23, 322–336.

Starkey, M.L., Bartus, K., Barritt, A.W., Bradbury, E.J., 2012. Chondroitinase ABC promotes compensatory sprouting of the intact corticospinal tract and recovery of forelimb function following unilateral pyramidotomy in adult mice. The European journal of neuroscience 36, 3665–3678.

Tauchi, R., Imagama, S., Natori, T., Ohgomori, T., Muramoto, A., Shinjo, R., Matsuyama, Y., Ishiguro, N., Kadomatsu, K., 2012. The endogenous proteoglycan-degrading enzyme ADAMTS-4 promotes functional recovery after spinal cord injury. J Neuroinflammation 9, 53.

Vandamme, C., Adjali, O., Mingozzi, F., 2017. Unraveling the Complex Story of Immune Responses to AAV Vectors Trial After Trial. Hum Gene Ther 28, 1061–1074.

Vogelaar, C.F., 2016. Extrinsic and intrinsic mechanisms of axon regeneration: the need for spinal cord injury treatment strategies to address both. Neural Regen Res 11, 572–574.

Weidner, N., Ner, A., Salimi, N., Tuszynski, M.H., 2001. Spontaneous corticospinal axonal plasticity and functional recovery after adult central nervous system injury. Proceedings of the National Academy of Sciences of the United States of America 98, 3513–3518.

Whishaw, I.Q., Gorny, B., Sarna, J., 1998. Paw and limb use in skilled and spontaneous reaching after pyramidal tract, red nucleus and combined lesions in the rat: behavioral and anatomical dissociations. Behav Brain Res 93, 167–183.

Yuan, W., Matthews, R.T., Sandy, J.D., Gottschall, P.E., 2002. Association between protease-specific proteolytic cleavage of brevican and synaptic loss in the dentate gyrus of kainate-treated rats. Neuroscience 114, 1091–1101.

