## Supplementary material for "Astrocyte-selective AAV-ADAMTS4 gene therapy combined with hindlimb rehabilitation promotes functional recovery after spinal cord injury"

**Polymerase chain reactions expansion of ADAMTS4 ORF**

The human ADAMTS4 cDNA was purchased from Origene (#RC09226, NM_005099) and used as a template for polymerase chain reactions for insertion into a therapeutic plasmid whereby forward and reverse primers were designed based on the fully sequenced open reading frame. The product of the polymerase chain reactions was subject to gel electrophoresis to check the correct size of the product. The PCR product was found to be approximately 2.5 kb: the predict weight of the ADMATS4 gene (Supplementary Fig 1 A). The gene was then isolated and ligated into the therapeutic expression plasmid: pAM/ITR-GfaABC_1_D-ADAMTS4-WPRE-BGHpA-ITR.

**Plasmid map of GfaABC_1_D-ADAMTS4 and restriction analysis**

The correct DNA sequence of the plasmid in relation to the plasmid mad constructed was confirmed using restriction enzyme digests (Supplementary Figs 1 A-C, 2). The same restriction enzymes used to insert the ADAMST4 gene were used; SpeI and HindIII. Two fragments were produced by this enzyme treatment. The sizes of the fragments produced by restriction enzyme digestion were consistent with the predicted fragment sizes calculated (5100 bp plasmid and 2387 bp transgene) therefore confirming the correct plasmid DNA sequences represented in the plasmid map. The restriction enzymes KpnI and BamHI were used to observe three fragments that correspond to the GfaABC_1_D, a 687 bp segment of the ADAMTS4 transgene and two fragments (1240 bp and 5560 bp) of the remaining 6.8 Kb or the plasmid. Following these confirmations the transgene was sequenced to confirm the correct DNA sequence was inserted into the plasmid. This sequencing was conduction by Massey Genome Services (Massey University) using appropriate forward and reverse primers. Sequencing data revealed that the ADAMTS4 gene sequence in the plasmid matched 100% to the gene sequence of the human ADAMTS4 gene. The expression plasmid was then packaged into an AAV5 vector. The vector was titre-matched to control for variability between tittering runs. To confirm the presence and purity of the vector stocks, a sample of each vector was subject to SDS-PAGE and then stained with coomassie blue to visualise the presence of the AAV capsid proteins; VP1 (87 KDa), VP2 (73 KDa) and VP3 (62 KDa). The AAV5-GfaABC_1_D-ADAMTS4 packaged vector (from now referred to simply as AAV-ADAMTS4) showed these three proteins at the correct ratios and only minor levels of contaminant were detected on the gel (Supplementary Fig. 1 C).

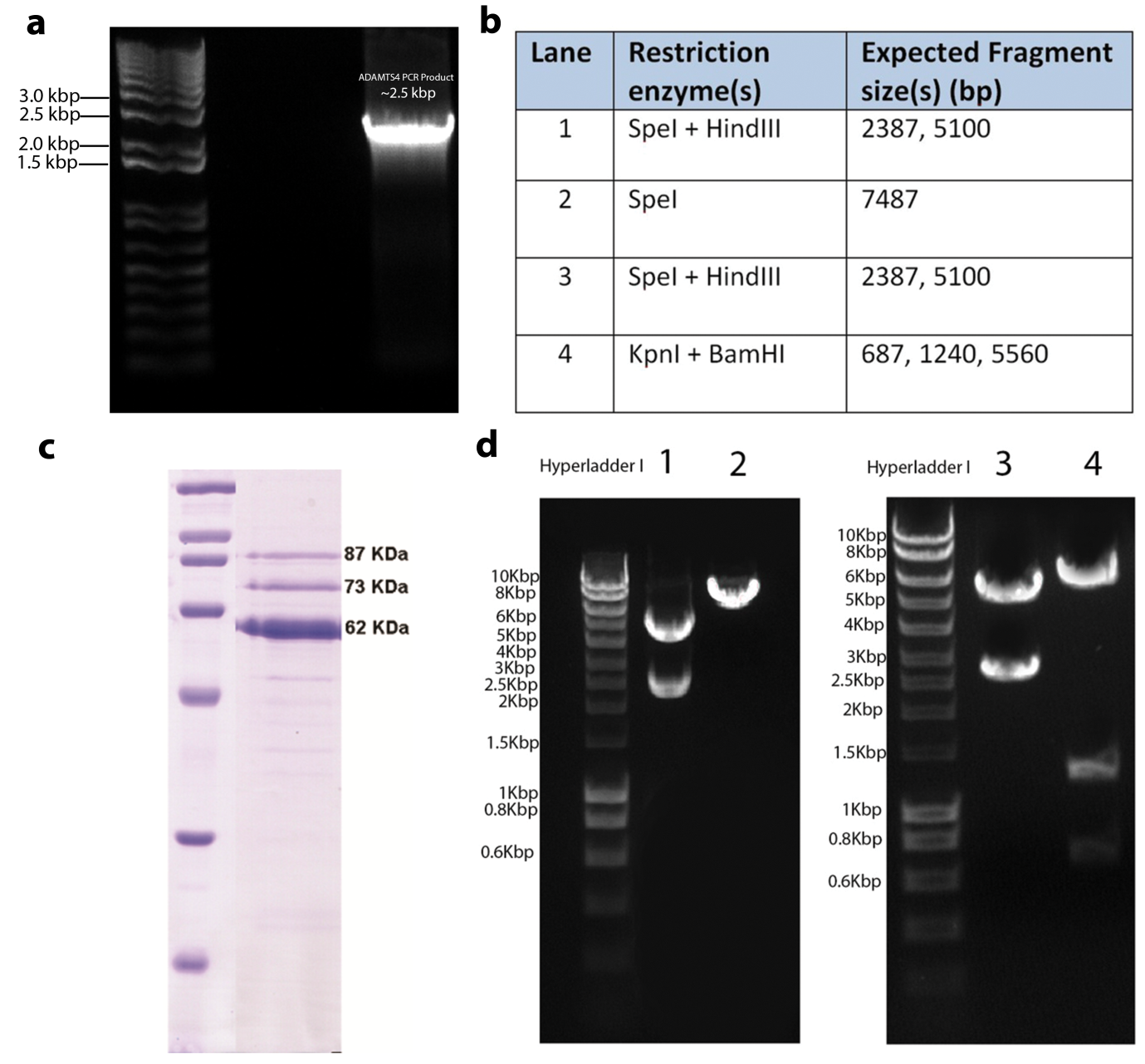

**Supplementary Figure 1: PCR product**. Primers specific to the human ADAMTS4 gene were designed and used for PCR to amplify the ADAMTS4 ORF. Agarose gel electrophoresis was used to confirm the correct size of the product before being isolated from the gel (a). Restriction enzyme analysis confirmed the correct fragments of the plasmid (b, d). SDS-PAGE confirmed that the vector stock produced was of sufficient purity (c).

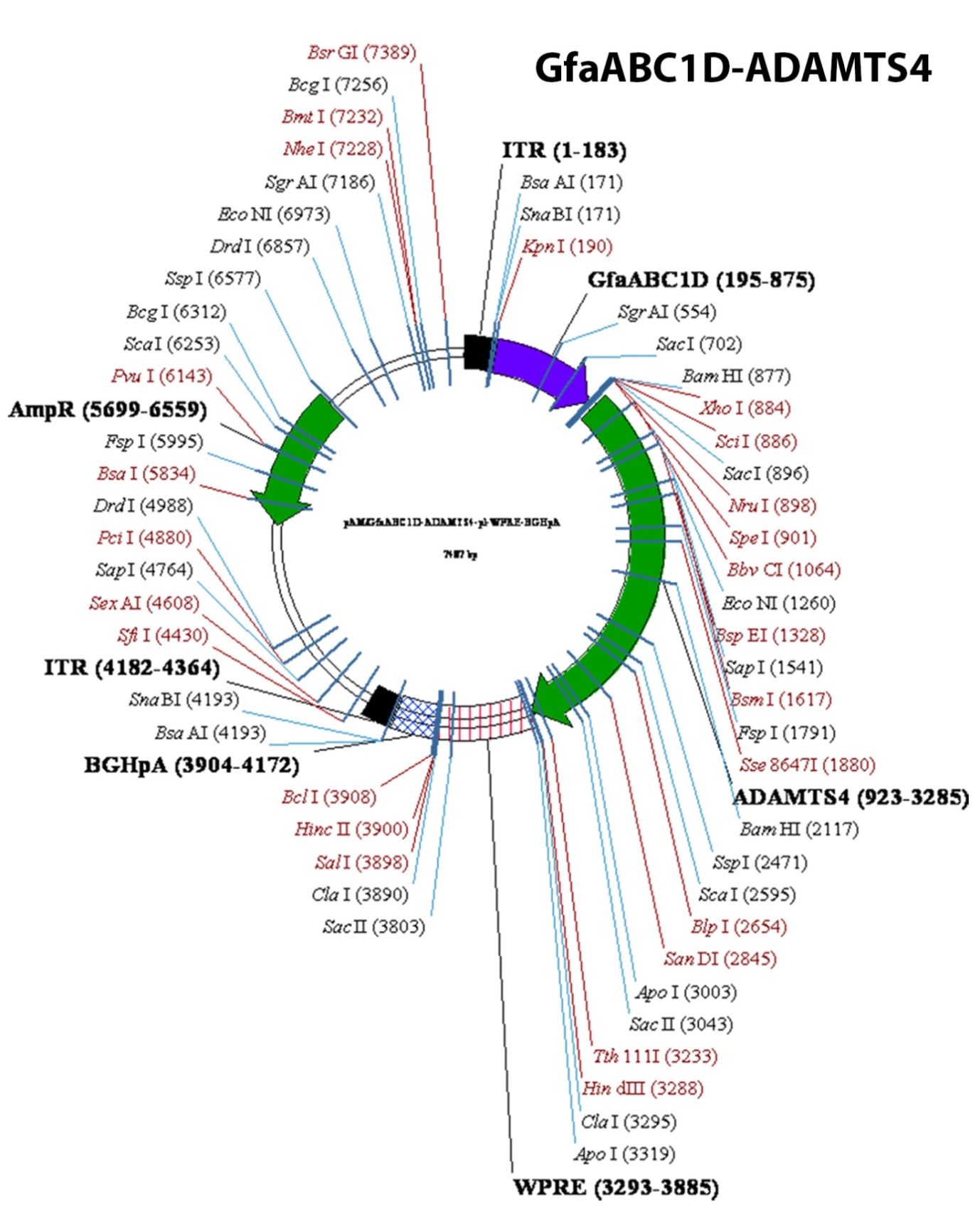

**Supplementary Figure 2: The plasmid map of AAV5-GfaABC_1_D-ADAMTS4**.

The endogenous proteoglycan degrading enzyme ADAMTS4 was packaged into an AAV expression cassette under the control of the truncated GfaABC_1_D promoter.

**SDS-PAGE to determine vector purity**

Coomassie Blue-staining of sodium dodecyl sulphate polyacrylamide gel electrophoresis (SDS-PAGE) was used to determine the purity of AAV vector stocks (Supplementary Fig 1 C). The separating gel consisting of 12% (w/v) acrylamide (BioRad, Hercules, CA) in 375mM Tris-HCl, pH 8.8, containing 0.1% (w/v) SDS was prepared, and polymerisation was catalysed by the addition of 0.03% tetramethylethylenediamine (TEMED, BioRad) and 0.08% ammonium persulphate (BioRad). The separating gel was poured into a gel casting chamber and overlaid with 500 μL of 50% isopropanol to level the interface between the separating and the stacking gels. Once the separating gel had set, the isopropanol was decanted. The stacking gel consisting of 5% (w/v) acrylamide, 0.13% bis-acrylamide, 125 mM Tris pH 6.8, 0.1% (w/v) SDS, 0.1% (w/v) ammonium persulphate and 0.1% (v/v) TEMED was prepared and poured above the separating gel. A comb was inserted to form wells for samples. 10 μL of sample buffer (0.2 M Tris-HCl, pH 6.8, 30% glycerol, 2% SDS, 0.03% bromophenol blue and 4% 2-mercaptoethanol) was added to 10 μL aliquot of each AAV vector preparation and mixtures were heated at 95ºC for 10 minutes to denature the AAV capsid proteins. The heat-denatured samples were loaded into wells and the proteins were separated by electrophoresis using 180 V current in an electrophoresis tank buffer (25 mM Tris, 192 mM glycine, 0.1% (w/v) SDS) until the bromophenol blue dye front reached the bottom of the gel. A protein molecular weight standard (Broad Range, Bio Rad) was also loaded on the gel to allow for determination of protein size. Following electrophoresis, the running gel was removed from the gel plates and fixed in 100 mL of fixative solution (50% (v/v) methanol, 10% (v/v) acetic acid) for 30 minutes. To visualise the protein bands, the gel was stained with Coomassie blue stain (50% (v/v) methanol, 0.05% (v/v) Coomassie Brilliant Blue R-250 (BioRad), 10% (v/v) acetic acid) for two hours. The gel was destained in 5% (v/v) methanol, 7% acetic acid overnight to remove non-specific background staining.

**Chondroitin sulphate proteoglycans are a substrate for ADAMTS4-mediated degradation**

Sodium dodecyl sulphate polyacrylamide gel electrophoresis (SDS-PAGE) and western blotting was used to validate that CSPGs are a substrate for ADAMST4 degradation, as well as detecting ADAMTS4 transgene expression in transduced astrocyte cultures. A vial of chick brain CSPGs (Merck Millipore) was thawed and 10 µg of protein was added to two 0.5 mL tubes. Recombinant human ADAMTS4 was then added to produce a concentration of 1 µM, diluted in 50 mM HEPES, 50 mM NaCl2, 10 mM CaCl2 at pH 7.5. The proteoglycans and ADAMTS4 were then incubated for three hours at 37°C. 6x Laemmli loading buffer was then added to each sample and they were loaded onto a 4-15% acrylamide gel and run at 40V for 10 mins and then 120V for 40 minutes. The gel was removed from the apparatus and fixed with 50% methanol, 10% acetic acid, 40% water for 30 minutes and then stained with the same solution plus 0.25% coomassie blue R-250 overnight on a rocker. The gel was then de-stained with 67.5% water, 7.5% acetic acid and 25% methanol for several hours until the background was clear. Un-digested proteoglycans are too large (<400kDa) to run on a gel, however, the digested forms of proteoglycans can. Supplementary Fig. 3 demonstrates that chondroitin sulphate proteoglycans cannot migrate in SDS-PAGE. However, shown in lane C, proteoglycans that had been incubated with ADAMTS4 are degraded and are able to separate on the gel. The molecular weight of ADAMTS4 is 53kD and therefore the band corresponding to that molecular weight in lane C is likely to be the ADAMTS4 enzyme.

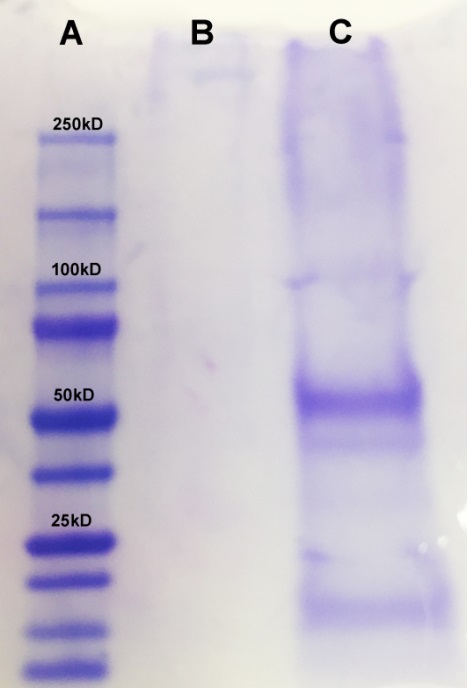

**Supplementary Figure 3: Chondroitin Sulphate Proteoglycans are a substrate for ADAMTS4-mediated degradation.**

10 µg of CSPGs isolated from chick brain were incubated with 1 µM ADAMTS4 at 37°C for three hours before being subject to SDS-PAGE and Coomassie blue staining. A) ladder; B) undigested CSPGs; C) CSPGs after incubation with ADAMTS4 showing digested fragments of CSPGs on the gel.

**Neuron cell culture**

All animal procedures were approved by the University of Auckland Animal Ethics Committee and performed in accordance with the New Zealand Animal Welfare Act 1999. Glass cover-slips were coated in 10 µg/mL poly-d-lysine (PDL) overnight at room temperature before being washed twice with warm sterile water. Pure extracellular CSPGs (Merck Millipore; #CC117) were used to coat the PDL coverslips by diluting them to a concentration of 20 µg/mL and incubating at 37°C for three hours. The proteoglycans were digested using recombinant human ADAMTS4 (R&D systems; #4307-AD-020). The enzyme was diluted to 20 nM in 50 mM 4-(2-hydroxyethyl)-1-piperazineethanesulfonic acid, 50 mM NaCl2, 10 mM CaCl2 and pH 7.5 and incubated for three hours at 37°C before being washed with sterile PBS. Isolation and culture of primary cortical neurons were performed as previously described^43^. Cells were maintained in a 37°C, 5% CO2 incubator (HeraCell; Thermo Fisher Scientific). An initial full medium change was performed 24 h after plating, followed by a 50% medium change twice weekly. Three days after plating the cells were fixed with 4% PFA-PBS at 4°C for 10 minutes. The neurons were cultured for two days before being processed for ICC to detect the neuronal marker β-tubulin 3. Neurons plated on CSPG substratum resulted in decreased neurite numbers and decreased neurite length compared to PDL alone (Supplementary Fig. 4 A). ADAMTS4 digestion of CSPGs from coated coverslips reversed the inhibitory effects of CSPG on neurite number and length (Supplementary Fig 4 B, C). No changes to the number of nuclei resulted from treatment with CSPGs indicating that the changes to neurite numbers and neurite length are unrelated to cell viability (Supplementary Fig. 4 D).

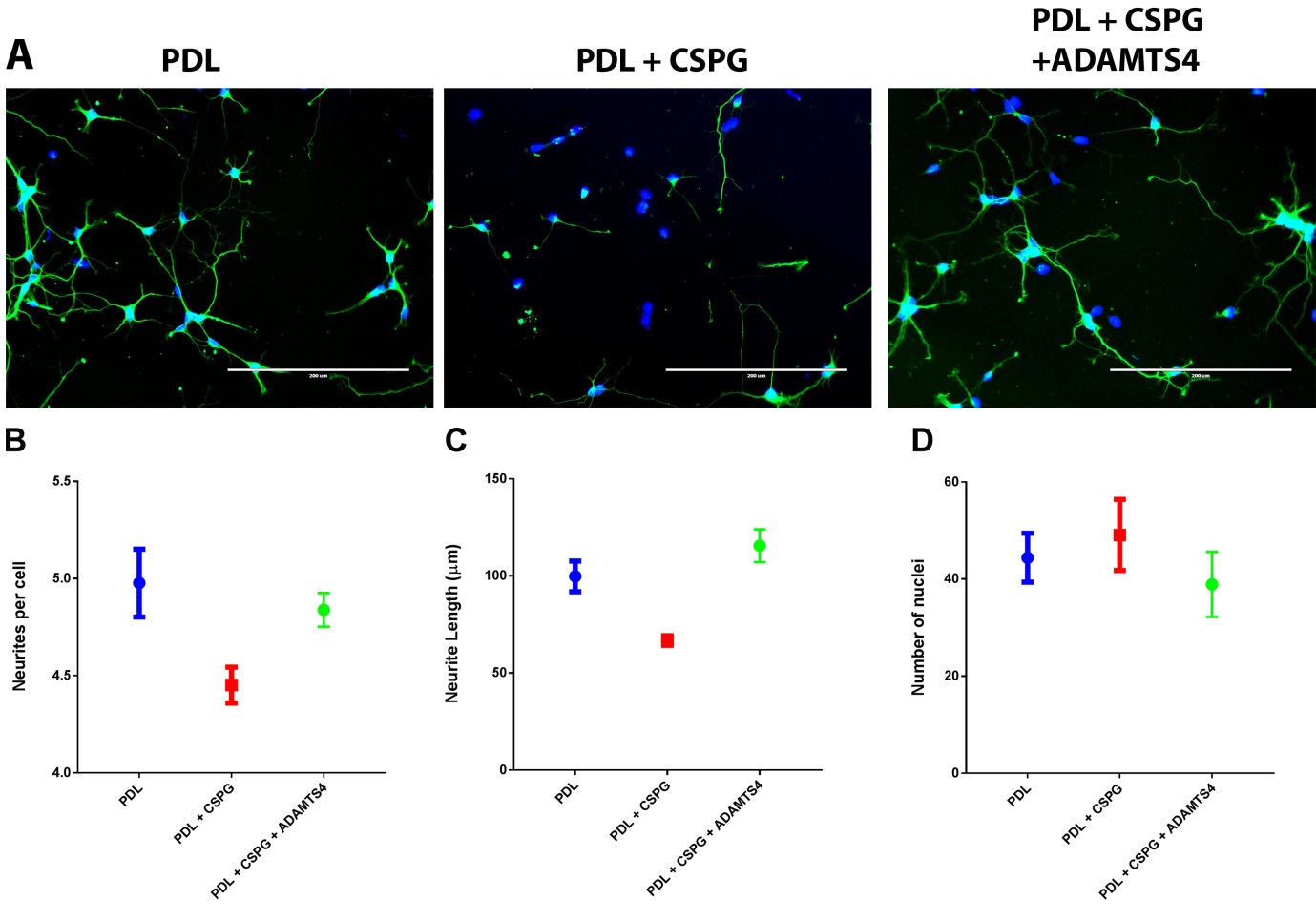

**Supplementary Figure 4: ADAMTS4 reverses CSPG-induced inhibition of neurite outgrowth.**

E16 cortical neurons were cultured on PDL, PDL+CSPG, or PDL+CSPG treated with ADAMTS4. Images were captured at 20x magnification on an EVOS FL Auto microscope and representative images are displayed (A). The number of neurites per cell (B) and neurite lengths (C) were counted from over 100 individual neurons and graphed. Data represent the mean ± SEM (n = > 100 cells for each group).

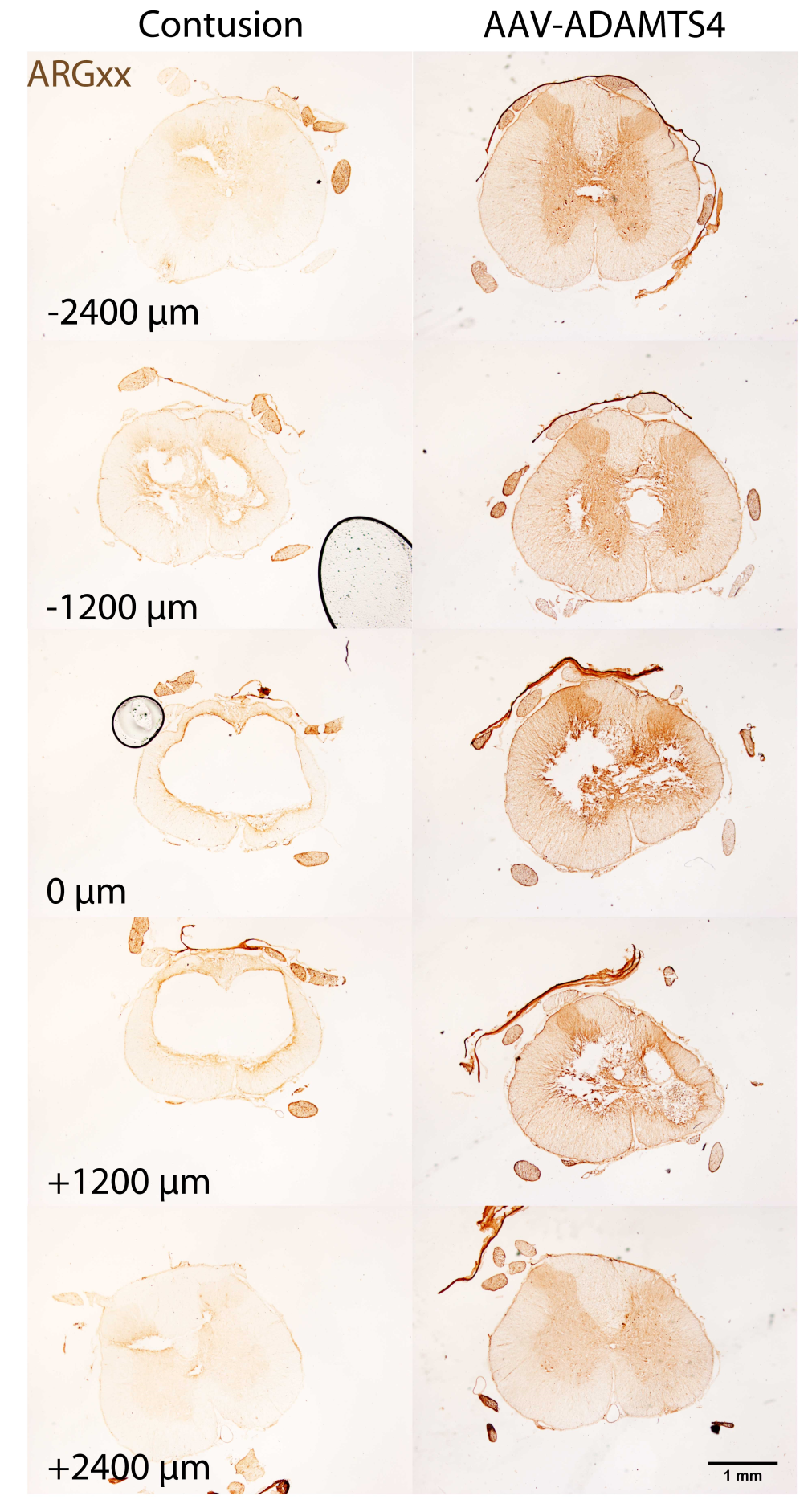

**Supplementary Figure 5: A neo-epitope produced by AAV-ADAMTS4 is evident throughout a long distance of spinal cord tissue.** Immunohistochemistry to detect the ADAMTS4-specific neoepitope ARGxx was used as a proxy for ADAMTS4 proteolysis. The neoepitope was abundant an expressed through more than 4800 µm of tissue. Scale bar = 1mm.
